# Reproductive modes in populations of late-acting self-incompatible and self-compatible polyploid *Ludwigia grandiflora* subsp. *hexapetala* in western Europe

**DOI:** 10.1101/2024.03.21.586104

**Authors:** Solenn Stoeckel, Ronan Becheler, Luis Portillo-Lemus, Marilyne Harang, Anne-Laure Besnard, Gilles Lassalle, Romain Causse-Védrines, Sophie Michon-Coudouel, Daniel J. Park, Bernard J. Pope, Eric J. Petit, Dominique Barloy

## Abstract

Reproductive mode, *i.e.,* the proportion of individuals produced by clonality, selfing and outcrossing in populations, determines how hereditary material is transmitted through generations. It shapes genetic diversity and its structure over time and space, which can be used to infer reproductive modes.

*Ludwigia grandiflora* subsp. *hexapetala* (*Lgh*) is a partially clonal, polyploid, hermaphroditic, and heteromorphic plant that recently colonized multiple countries worldwide. In western Europe, individuals are either self-incompatible caused by a late-acting self-incompatibility (LSI) system developing long-styled flowers, or self-compatible (SC), with short-styled flowers.

In this study, we genotyped 53 long- and short-styled populations newly colonizing France and northern Spain using SNPs to estimate rates of clonality, selfing and outcrossing. We found that populations reproduced mainly clonally but with a high diversity of genotypes along with rates of sexuality ranging from 10% up to 40%. We also found evidence for local admixture between long- and short-styled populations in a background of genetic structure between floral morphs that was twice the level found within morphs. Long- and short-styled populations showed similar rates of clonality but short-styled populations presented significantly higher rates of selfing, as expected considering their breeding system, and despite the small rates of failure of the LSI system. Within the 53 studied populations, the 13 short-styled populations had fewer effective alleles, lower observed heterozygosity, and higher inbreeding coefficients, linkage disequilibrium and estimates of selfing than what was found in long-styled populations. These results emphasize the necessity to consider the variation of reproductive modes when managing invasive plant species. The overall maintenance of higher genetic diversity with the possibility of maintaining populations clonally in the absence of compatible partners may explain why long-styled individuals seem to be more prevalent in all newly expanding populations worldwide. Beyond *Lgh*, our methodological approach may inspire future studies to assess the reproductive modes in other autopolyploid populations.

## Introduction

Plants reproduce using different mechanisms (i.e., breeding system), including clonality and different types of sexuality (i.e., mating system, including allogamy and selfing; Holsinger 2000). Reproductive mode corresponds to the actual balance between sexual and clonal reproduction, measured by the rate of clonality (De Meeus et al. 2007; Stoeckel et al. 2021a), and, within the proportion of offspring produced by sexuality, the balance between autogamy and geitonogamy (hereafter *selfing*), and allogamy (*outcrossing*), measured by the rate of selfing (Bürkli et al. 2017). Reproductive mode is one of the main drivers of genetic diversity in populations and species (Duminil et al. 2007, Ellegren & Galtier 2016, Glémin et al. 2019). It determines the way hereditary material is transmitted over generations, and thus constrains the range of genetic diversity that can evolve within populations and species (De Meeus et al. 2007, Fehrer 2010, Stoeckel & Masson 2014, Rouger et al. 2016). Its genetic consequences are deep enough to be discernible even in the presence of other affecting factors (Charlesworth 2003). Considering its influence on population genetic diversity and structure, reproductive mode is one of the major biological traits to decipher before interpreting other biological and ecological forces using molecular data (Duminil et al. 2007, Reichel et al. 2016, Bürkli et al. 2017, Orive & Krueger-Hadfield 2021). Given an adequate theoretical framework, population geneticists can use genotypic data to infer reproductive modes in populations (Halkett et al. 2005, Arnaud-Haond et al. 2007, Hardy 2016, Bürkli et al. 2017, Stoeckel et al. 2021a).

Many plant species reproduce using different breeding systems they adapt to different ecological contexts resulting into different reproductive modes (Richards 1997, Charlesworth 2006). Uniparental reproduction, including clonality and selfing, may help plants to spread into new areas (Baker’s conjecture: Barrett et al. 2008, Pannell et al. 2015). Clonality consists of a parent producing a new descendant that is a genetic copy of itself with the exception of rare somatic mutations and mitotic recombinations (De Meeus et al. 2007). It results into an average excess of heterozygotes compared to Hardy-Weinberg expectations (*i.e.,* negative F_IS_), large variance of F_IS_ along the genome and a small excess of linkage disequilibrium, increasing with decreasing population size (Navascues et al. 2010, Stoeckel & Masson 2014, Stoeckel et al. 2021a). When parents can yield multiple descendants by clonality, it may generate repeated genotypes in populations (Arnaud-Haond et al. 2007). These effects vary with the rates of clonality and, which in turn, can be used to infer these rates within populations (De Meeus et al. 2006, Becheler et al. 2020, Arnaud-Haond et al. 2020). Hermaphroditic plants have the possibility to sexually reproduce with themselves (selfing). Selfing decreases effective population sizes and gene diversity resulting into high probability of allele fixing (Wright et al. 2008, Roze 2016, Glémin et al. 2019). Selfing also prohibits genetic mixing between different ancestral lineages, which decreases heterozygosity within individuals (David et al. 2007, Hardy 2016) and strongly increases linkage disequilibrium between genes along genomes (Golding & Strobeck 1980, Norborg 2000, Lucek & Willi 2021). Around half of hermaphrodite plants restrict self-fertilization using a variety of molecular mechanisms grouped under the term “self-incompatibility (SI) systems” which favour outcrossing within populations (De Nettancourt 2001, Castric & Vekemans 2004, Gibbs 2014, Steinecke et al. 2022). Outcrossing is overall expected to increase genetic diversity, to limit allele fixation and to reduce linkage disequilibrium when compared to selfing, but only if linked to the genes involved in self-recognition (Glémin et al. 2001, Stoeckel et al. 2006, Navascues et al. 2010). Within the different identified SI systems (Charlesworth et al. 2005, Franklin-Tong 2008), the late-acting SI system (hereafter LSI) is still poorly studied despite being identified in multiple taxonomic groups in angiosperms (De Nettancourt 1997, Gibbs 2014). In LSI systems, self-pollen tubes grow and are only blocked shortly before penetrating the ovule. Such species may present reduced female fertility due to self-pollen disabling ovules, favouring the clonal regeneration of populations (Vaughton et al. 2010). LSI systems are also characterized by low but recurrent failures of the self-recognition system due to its late mechanism, leading to the production of a low amount of selfed seeds in populations (Seavey & Bawa 1986; Chen et al. 2012, Gibbs 2014). In contrast, gametophytic and sporophytic self-recognition occurring very early in the pistil drastically limit self-pollination and thus selfed seeds (Lawrence 1985, Gibbs 2014). However, we do not know yet if these rare selfed seeds contribute to the dynamics of genetic diversity of LSI populations and species, especially in peripatric conditions or in any situation when compatible partners may lack. Due to these selfed seeds, LSI system may be relatively ineffective at driving genetic diversity within populations, contrary to gametophytic and sporophytic SI systems (Brennan et al. 2002, Stoeckel et al. 2006, Koelling et al. 2011, Busch & Urban 2011). In such situations, the emergence and maintenance of LSI systems appear as new evolutionary puzzles among the reproductive systems.

Water primrose*, Ludwigia grandiflora* subsp. *hexapetala* (Hook. & Arn.) Nesom and Kartesz (2000), hereafter *Lgh*, is an insect pollinated, partially clonal, hermaphroditic and heteromorphic plant supposed to be native from central and south America. This species is one of the most invasive aquatic plants in the world (Thouvenot et al. 2013). *Lgh* is a decaploid species (2n=10x=80 chromosomes), resulting from hybridization of different ancestral diploid species, some of which are represented more than once in the total genome of *Lgh*, which belongs to the genus *Ludwigia* L. section Jussiaea (Hoch et al. 2015, Barloy et al, 2024). Interestingly, *Lgh* includes an autotetraploid set of chromosomes, shared with *L. peploides* subsp. *montevidensis* (2n=2x=16), hereafter *Lpm*, that is clearly distinct from the other ancestral part of the *Lgh* genome (Barloy et al. 2024). *Lpm* is a self-compatible-only diploid species with only one common floral morphology (Estes & Thorp 1974, Grewel et al. 2016). *Lgh* presents heteromorphic flowers corresponding to two floral morphs: L-morph individuals develop long-styled flowers and S-morph individuals develop short-styled flowers, that cross and result into 100% viable and fertile F1 and F2 descendants (Portillo-Lemus 2021, Portillo-Lemus et al. 2021) while inter-species crosses only result in a low number of chlorotic and unfertile descendants (Barloy et al. 2024).

All tested L-morph flowers expressed an active LSI in western European populations, with only one self-incompatible type detected so far (Portillo-Lemus et al. 2022). During the core flowering season (summer), in experimental greenhouse conditions, L-morph individuals show a stable seemingly insignificant rate of autogamy (around 0.2‰ of the available ovules) that increases at the end of the flowering season, during autumn, to 1‰, which is common in LSI systems (Gibbs 2014). Due to the massive blossoming of this species, growing in very dense populations, these selfed seeds would add up yearly in field populations to a hundred seeds per square meter (Portillo-Lemus et al. 2022). This pattern of low rates of autogamy, that increases at the end of the flowering season, may provide the advantage of reproductive assurance (Goodwillie & Weber 2018). In contrast, all tested S-morph individuals were self-compatible (SC) in western European populations but in their pistils, pollen tubes of the L-morph growed significantly faster and were thus advantaged to fertilize ovules when in competition with self-pollen (Portillo-Lemus et al. 2022). In addition, peripatric *Lgh* populations, including European populations, were previously reported as exclusively clonal with few clonal lines (Dandelot 2004, Okada et al. 2009). Recently established populations in France and northern Spain mostly present only one of the two compatibility modes locally, sometimes with a population of the other type a few to tens of kilometers away, which may result in effective allogamy (Portillo-Lemus et al. 2021, 2022). All these different breeding systems make possible very different reproductive modes in populations, comprising all possible quantitative combinations of mainly clonal, autogamous and allogamous modes.

Here, we assessed the reproductive modes of 53 invasive *Lgh* populations across western Europe. Considering the complex case of *Lgh* in western European populations, we hypothesized that L-morph populations supposed to express a LSI should present typical genetic footprints of dominant outcrossing, or alternatively higher rates of clonality due to the local lack of compatible partners and self-pollen interferences, while S-morph populations supposed to be SC should present higher rates of sexuality prevailed by selfing. To understand the genetic impacts of reproductive modes, we also quantified the covariations of genetic indices and their importance to define the genetic diversity within the 53 genotyped populations, and compared these observations to the theoretical expectations obtained from a Wright-Fisher-like model extended to autotetraploids (Stoeckel et al. 2024). Finally, we took advantage from the fact that recent local populations in France and northern Spain still present only one of the two compatibility modes to compare and interpret the genetic differences found within and between L-morph (LSI) and S-morph (SC) populations, with the aim to assess the influence of LSI on genetic diversity, as similarly tackled in sporophytic self-incompatible and SC Brassicaceae species previously (Koelling et al. 2011).

## Materials and methods

### Sampling strategy and floral morphology of populations

We collected 795 stems of *Lgh* from 53 locations (52 locations in France and one in Catalonia, Spain), corresponding to an area that spans 580 kilometres east-to-west and 1,100 kilometres north-to-south (Figure SI1). At each location, we collected 15 stems (hereafter, ‘individuals’) along a linear transect of 40 meters. Along each transect, we randomly collected three stems at coordinates X_1_ = 0m; X_2_ = 10m; X_3_ = 20m; X_4_ = 30m and X_5_ = 40m within a one meter-square quadrat. The young leaves of each sampled individual were stored after lyophilization until DNA extraction.

We visually identified floral morphologies of flowers found along each transect within the sampled *Lgh* populations. In a previous study on seven populations among the 53 studied here (underlined population names in Figure SI1), one hundred and five sampled individuals resulted to identify a binary distribution of floral morphology with formally-identified self-incompatibility types: all L-morph individuals were LSI typed and all S-morph individuals were SC typed (Portillo-Lemus et al 2021) while all these individuals succeeded to cross and give viable and fertile plants. Interestingly, these two types of populations spatially distribute in monomorphic populations along different rivers. We supposed for this study the LSI versus SC status of individuals and populations using their floral morphologies: L-morph individuals were supposed to develop a LSI system with only one self-incompatible type found so far and S-morph individuals being SC. To support this conjecture, as done in Portillo-Lemus et al. (2021), we checked the fruitset in each of the 53 sampled and genotyped populations: A low fruitset, or even no fruit at all were found in L-morph individuals and populations, while full fruitsets were found in S-morph individuals and populations, in agreement with our conjecture.

In addition to the crossbreeding results in which all L- and S-morph individuals succeeded to cross, giving full fruit set and 100% viable first- and second-generation descendants (Portillo-Lemus et al. 2021), we here counted the chromosome numbers on karyotypes of S-morph individuals sampled in two fruitful populations and of L-morph individuals sampled in five fruitless populations to validate that L- and S-morph individuals belong to *Lgh*. Between 50 to 150 kilometers separated two consecutive samples (populations underlined Figure SI1). To prepare the karyotypes, we used the method detailed in Barloy et al (2024) that already karyotyped a S-morph individual sampled near the French Atlantic coast. The same method was used in Bou Manobens et al. (2019) to karyotype a L-morph individual sampled in Catalunya.

### Definition of the autotetraploid SNP marker set

As no molecular markers suitable for clonal discrimination were yet available for *Lgh* and *Lpm*, we generated an original set of SNP markers via RAD-Seq (Baird et al., 2008) using two pools of 15 individuals each, respectively sampled across five *Lgh* and three *Lpm* western European populations. RAD DNA library generation, sequencing and the analysis pipeline to identify *Lgh* SNP markers were carried out as described in Delord et al. (2018). In brief, DNA of *Lgh* and *Lpm* were digested by *Sbfl* restriction enzyme and used to prepare DNA libraries that were then sequenced using an Illumina HiSeq3000 (150bp paired-end reads). A total of 14,233 and 34,287 RAD-Seq-determined SNPs were filtered to yield 340 and 326 SNPs, one per aligned sequence, for *Lpm* and *Lgh*, respectively. Design of primers compatible with Hi-Plex multiplexing was carried out by Melbourne Bioinformatics. Finally, sixty and fifty SNP markers matched the quality criteria to design a Hi-Plex set of SNPs for *Lpm* and *Lgh*, respectively (Hammet et al. 2019). In our study, we only considered polymorphic SNPs that were shared between *Lgh* and *Lpm* to be sure they belong to the tetraploid part of *Lgh* genome derived from *Lpm* (Barloy et al. 2024). We finally kept this set of 36 polymorphic and stable SNP markers to genotype *Lgh* samples and analyze genetic diversity in each sampled population (primer sequences are openly listed in Stoeckel et al. 2023).

To verify that this set of SNPs was really tetraploid, we computed the Akaike’s information criterion from the maximum likelihood of the best genotype considering the distribution of sequenced allele countings among individuals and markers as a function of the ploidy level, using a similar approach to that proposed by Burnham & Anderson (2002).

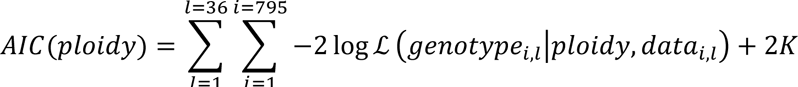

where *data_i,l_* is the distribution of sequenced alleles among A, C, G and T within individual *i* at locus *l*, and K is the number of possible genotypes given the ploidy and the four possible alleles (A, C, G and T). The likelihood of the possible genotypes ℒ(*genotype_i,l_*|*ploidy, data_i,l_*) follows a multinomial distribution of the sequenced allele countings distributed between the four different possible nucleobases (A, C, G and T).

### DNA extraction, SNP marker production and genotyping

To genotype the 795 individually-tagged samples with these 36 SNPs, we extracted DNA from 25 to 30 mg of dried young leaf tissues. The DNA extractions were performed using the NucleoSpin Plant II from MACHEREY-NAGEL kit, following the manufacturer’s recommendations. We used L1 buffer as lysis buffer. All individuals were genotyped from a solution of 20ng/μl of *Lgh* DNA, using a modified version of the Hi-Plex protocol (Hammet et al. 2019; Besnard et al. 2023). Hi-Plex is an amplicon sequencing technique (*sensu* Meek & Larson 2019) in which all SNPs are co-amplified in a multiplex reaction before Illumina or Ion-Torrent sequencing. Here, we used Illumina. Intermediate steps include dual indexing of individual samples used for demultiplexing. Reads were then assigned to loci by aligning them to reference sequences with BWA-MEM 0.7.15-r1140 (Li & Durbin 2010) and alleles were counted with Samtools 1.9 (Danecek et al. 2021).

### Allele dosage

The posterior probabilities of each single SNP genotype within each individual (hereafter *single SNP genotype*) were computed using the likelihood of all possible genotypes considering a multinomial distribution of the number of times each nucleobase was genotyped at one SNP marker within one individual. In order to obtain the most confident dataset possible, we only assigned a genotype when its posterior probability of allele dosage exceeded 70%. When one SNP within one individual presented a posterior probability of allele dosage equal or lower than 70%, we assigned and analysed it afterward as a missing genotype.

### Genotypic and genetic descriptors

We expected that populations reproducing clonally may yield repeated multi-locus genotypes (MLGs, i.e., the same genotype at all the 36 SNPs found in multiple individuals). By possibly producing these repeated genotypes and by varying the relative distribution of the number of samples of each of these distinct genotypes, rates of clonality impact genotypic richness and evenness in populations (Halkett et al. 2005; Arnaud-Haond et al. 2007).

We measured genotypic richness using the R index (Dorken & Eckert 2001, Arnaud-Haond et al. 2005), which is defined as:

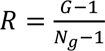

where G is the number of distinct genotypes (genets) and N_g_ is the number of genotyped individuals. We also measured genotypic evenness as the parameter β of the Pareto distribution, which describes the slope of the power-law inverse cumulative distribution of the size of MLGs (Arnaud-Haond et al. 2007):

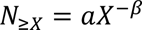

where N_≥*X*_ is the number of ramets in genets containing *X* or more ramets in the sampled population, and the parameters *a* and *β* are fitted by regression analysis. Low β-values indicate the dominance of one or few clonal lineages. Pareto β is less biased by the sampling effort than clonal richness R (Stoeckel et al. 2021a). In our sample of 15 individuals, we expected a population with only one repeated MLG to present a Pareto β value of 0.03 and a value of 3 in samples with no repeated genotypes. β < 2 indicates a population reproducing with high rates of clonality (greater than 0.8 to 0.9). For a sample size of 15 individuals, 2 < β < 3 indicates intermediate to low rates of clonality (0.6 to 0.8). A β at 3, its maximum value, is indicative of a mainly sexual population, with rates of clonality ranging from zero to 0.6, depending on the other genetic indices.

Rates of clonality and selfing confer different effects on the range of within-population genetic polymorphism as well as on probabilities of genetic identities within individuals. We thus also estimated expected and observed heterozygosity H_E_ and H_O_, allelic richness A_E_, inbreeding coefficient F_IS_, and linkage disequilibrium within each sampled population.

The inbreeding coefficient F_IS_ (Wright 1931, 1949) accounts for intraindividual genetic variation at one locus as a departure from Hardy-Weinberg assumptions of the genotyped populations. We computed one F_IS_ value per locus per population. We then reported the mean values of F_IS_ (MF_IS_) and the inter-locus variance of F_IS_ (VarF_IS_) for each population. Both MF_IS_ and VarF_IS_ are very informative about the underlying reproductive systems (Stoeckel et al. 2006, David et al. 2007). Clonality makes the MF_IS_ values tend toward −1 along with high interlocus variance (Stoeckel & Masson 2014, Stoeckel et al. 2021a). A moderate amount of sexual reproduction results in MF_IS_ values around 0 (Balloux et al. 2003). VarF_IS_ varies with rates of clonality, from very limited variance expected in sexual populations to high variance in very clonal populations (Stoeckel & Masson 2014, Stoeckel et al. 2021a). Positive MF_IS_ values are expected in populations reproducing using consanguinity and selfing (Castric et al. 2002, David et al. 2007). All sexual reproductive modes, allogamous and autogamous, result in low VarF_IS_ within a population as massive recombination tends to homogenize intra-individual genetic identities along the genomes (Stoeckel et al. 2021a). We measured linkage disequilibrium over all markers using the unbiased multilocus linkage disequilibrium index, *r̄_d_* (Agapow & Burt 2001). This mean correlation coefficient (*r*) of genetic distances (*d*) between unordered alleles at *n* loci ranges from 0 to 1. This metric has the advantage of limiting the dependency of the correlation coefficient on the number of alleles and loci, and it is well suited to measure linkage disequilibrium in partially clonal populations (De Meeus & Balloux 2004, De Meeus et al. 2006). In general, LD is only slightly affected by clonality, except when clonality is high (c > 0.9) and/or when genetic drift dominates over mutation rate (*e.g.,* when population sizes are small N<50 compared to u=0.001; Navascues et al. 2010, Stoeckel et al. 2021a). In contrast, inbreeding and selfing are efficient processes for quickly generating strong LD, after only few generations. Finally, we measured genetic differentiation between populations of LSI, between populations of SC and between LSI and SC populations using ρ_ST_, an index adapted to study autopolyploid populations (Ronfort et al. 1998). All these indices were computed using GENAPOPOP (Stoeckel et al. 2024), a software dedicated to analysing genetic diversity and differentiation in autopolyploid populations genotyped with confident allele dosage and reproducing through all possible rates of clonality and selfing.

We also used SPAGEDI (V1.5, Hardy & Vekemans 2002) to estimate rates of selfing within autopolyploid populations genotyped with confident allele dosage. This approach infers rate of selfing (Sg) from identity disequilibrium coefficients (g_2z_ estimator), assuming that populations reproduce by self-fertilization and random mating and are at inbreeding equilibrium (David et al. 2007 for diploids, Hardy 2016 for autopolyploids). Identity disequilibrium coefficients are measured as the correlation in heterozygosity of distinct loci within the genome, and present the advantage of being more robust to null alleles and genotyping errors than raw F_IS_ (David et al., 2007). The effect of partial clonality on g_2z_ estimator is not yet defined. We thus considered for this study that clonality would only marginally impact identity disequilibrium coefficients, the possibility to be at the inbreeding equilibrium and the corresponding estimates of selfing. A synthesis of all these hypotheses based on previous knowledge is provided in Table SI1.

### Ranking populations by clonality and selfing

We proposed and computed a synthesis index (Σclon) calculated from Pareto β and VarF_IS_ to rank the studied populations from the less to the more clonal. These two population genetic indices are known to vary with rates of clonality being unbiased in samples (Stoeckel et al. 2021a). To avoid issues of scaling as the range of values of these descriptors are different by several orders of magnitude, Σclon is computed in each population *i* as the sum of Pareto β(*i*) and VarFIS(*i*) values that were previously normalized over the whole dataset, respectively *Z*_β_(*i*) and *Z*_VarFIS_(*i*).

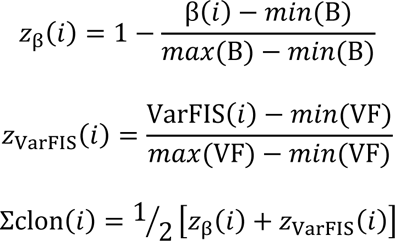

where β(*i*) and VarFIS(*i*) are the measured Pareto β and variance of F_IS_ over loci in the population *i*, and where Β = {*β*(1), *β*(2), …, *β*(55)} and *VF* = {VarFIS(1), VarFIS(2), …, VarFIS(55)} are the respective sets of Pareto β and VarF_IS_ over loci in the 53 populations used to obtain their minimum (*min*) and maximum (*max*) values.

Using the same approach as for Σclon, we computed a synthesis index (Σself) calculated from *r̄_d_*, MF_IS_ and Sg to rank the studied populations from the less to the more selfed.

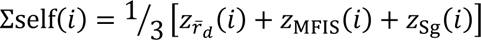

### Statistical data analysis

First, to better understand the correlation between population genetic indices and estimates of reproductive modes from a large field dataset and to compare with the theoretical correlations obtained from simulations (Stoeckel et al. 2024), we computed a Principal Component Analysis (PCA) to comprehend the covariations of genetic indices, the correlations and the redundancies between the 17 genetic diversity indices (described above: G, R, D, Pareto *β*, *r̄_d_*, Ae, varAe, He, varHe, Ho, varHo, MF_IS_, varF_IS_, PIDsib, PIDu, Sg, SE.Sg) measured among the 53 sampled populations, and their link with reproductive modes including self-compatibility. We reported the amount of variation retained by the first two principal components and the correlation circle on which we plotted predictions of Σclon and Σself as supplementary variables. We also reported the score plot to visualize how L-morph (LSI) and S-morph (SC) populations distribute along the principal components of population genetic diversity. To avoid scaling issues, all descriptors were normalized before analysis.

We then analyzed the relationship between genetic diversity indices and their relationships with Σclon and Σself measured in the 53 sampled populations using Kendal partial rank-order correlation tests. We reported the corresponding matrices of correlation.

To detect differences in distribution of population genetic indices, Σclon and Σself, among floral morphs, we computed non-parametric Kruskal-Wallis tests that do not make any assumptions about the type of distribution and about homogeneity of variances between the tested distributions. When needed, we used post-hoc pairwise tests for multiple comparisons of mean rank sums (Nemenyi’s test).

All statistical tests were computed using PYTHON 3.11, SCIPY.STATS 1.9.3 (Virtanen et al. 2020) and SCIKIT-POSTHOCS (Terpilowski 2019), except the PCA that was performed using R V4.2.2 and the library FACTOMINER (Lê et al. 2008).

## Results

Among the 53 populations, we found 40 populations with only L-morph individuals and 13 populations with only S-morph individuals (Table SI2, Figure SI1). Karyotypes of L-morph plants from two populations and of S-morph plants from five populations all presented the same number of chromosomes (2n=80, Figure SI2) confirming that L and S-morph individuals in France and L-morph individuals in northern Spain (Bou Manobens et al. 2019) belong to *L. grandiflora* subsp. *hexapetala* (2n=10X=80, Barloy et al. 2024). Akaike’s information criterion on the distribution of allele counting over all our data supported tetraploidy as the best ploidy level for this set of 36 SNPs (Figure SI3), as expected from Barloy et al (2024).

### Allele dosage and missing genotypes

Within the 795 individuals genotyped at 36 SNPs (resulting in a total of 28,620 single-SNP genotypes), 99.97% (28,612) of SNPs were genotyped with posterior probability of allele dosage superior to 70% (Table SI3). In total, 785 individuals were genotyped with a full set of 36 SNPs with posterior probabilities of allele dosage higher than 70%. Ten individuals distributed in nine populations showed one of their SNP markers with posterior probabilities of allele dosage equal or lower than 70%, that we assigned therefore as missing genotypes.

### Statistical power of the developed SNP marker set

Over the 36 polymorphic SNPs, we found an effective number of alleles (A_E_) of 1.36 per SNP over all populations (Table SI2). Among populations, mean A_E_ values over the 36 SNPs were homogenous, ranging from 1.22 to 1.55 (median=1.34). We, however, found large standard deviation of A_E_ between SNPs within populations, ranging from 1.6 to 2.3 (median=2.1). Some SNPs were apparently fixed in some populations while polymorphic at the scale of the whole dataset. When not fixed, gene diversity H_E_ in polymorphic SNPs ranged from 0.14 to 0.33 (median=0.2).

These 36 SNPs in the autotetraploid part of *Lgh* would theoretically allow 4^36^=4.7×10^21^ different possible MLGs considering the four possible nucleobases and 2^36^=6.9×10^10^ different possible MLGs assuming two possible nucleobases per locus. Considering allele frequencies in the sampled populations, the probabilities of identities under panmixia ranged from 8.5×10^-12^ to 5.2×10^-5^ (median=2.1×10^-7^) and the unbiased probabilities of identity between sibs P_ID-SIB_ ranged from 4.2×10^-6^ to 8.3×10^-3^ (median=5.6×10^-4^; Table SI2). We then considered that the SNP set we used to genotype the 795 sampled individuals via Hi-Plex method showed sufficient statistical power to distinguish between true MLGs, and that individuals with identical MLGs were true repeated genotypes (ramets) of a clonal lineage (a genet).

### Genetic and genotypic diversity

Across populations, we identified a total of 462 distinct MLGs (genets) within the 795 sampled individuals genotyped. Among them, we found 404 genets (88%) with a single ramet and only 58 genets (12%) with more than two ramets (Table SI4). Forty-eight genets had two to seven ramets distributed over one to six populations (median=2), seven genets with 10 to 33 ramets distributed over two to 17 populations (median=9), and one large genet of 99 ramets distributed over 24 populations (Figure 1).

**Figure 1:**
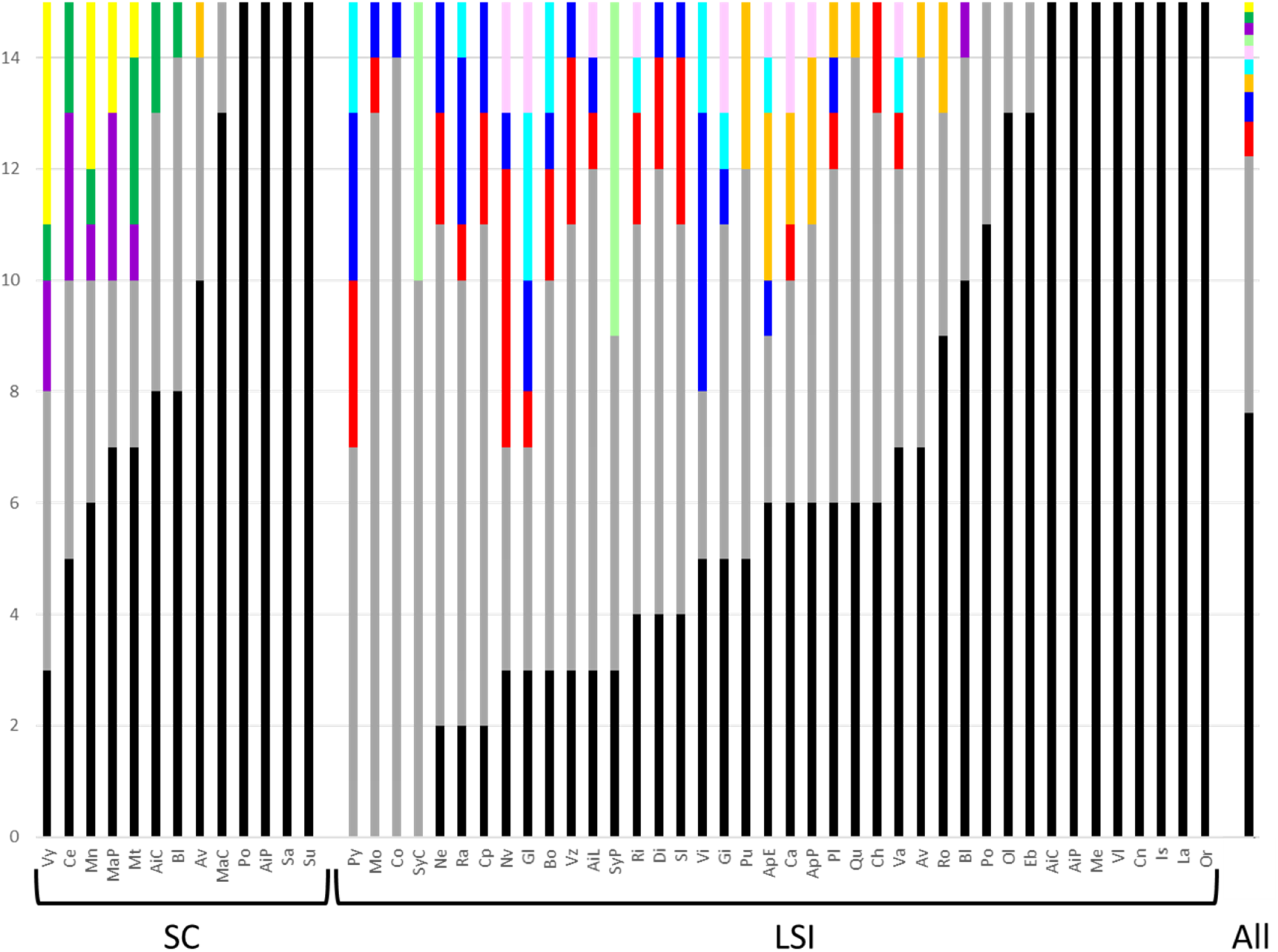
Distributions of Multi-Locus Genotypes (MLGs, we consider as genet) among LSI and SC *Lgh* populations in western Europe. The y-axis represents the distribution of the genotyped samples (ramets) with the different following categories: In black, the number of unique MLGs (MLGs that were found only once over all populations); In grey, the number of MLGs with less than 7 ramets over all populations; In color, MLGs with more than 10 ramets found in multiple populations (one color per different genet, identifying the same genet across all populations). Proportions of each types of MLGs over all the populations (795 genotyped individuals) are plotted in the last right bar (All). More than half the samples are unique genotypes (black bar).

Within populations, we found from three to 15 different genets (median=12) per population among the 15 sampled individuals (Table SI4), implying the clear occurrence of repeated genotypes but also a wide diversity of genets within most populations (Figure 1). Accordingly, genotypic richness (R) ranged from 0.14 to 1 (median=0.79). The genotypic evenness, Pareto β, ranged from 0.056 to 3 (median=1.478, Table SI2).

Observed heterozygosity H_O_ was also high in most populations, slightly above expected heterozygosity, ranging from 0.13 to 0.38 (median=0.26). The mean standard inbreeding coefficients (MF_IS_) averaged over all genotyped loci within populations were negative in all 53 populations and ranged from −0.33 to −0.126 with a very negative median of −0.274 (Table SI2). Variances of F_IS_ between loci within population were very high, ranging from 1.75 to 36.60 (median=27.57). All these measures argued for reproductive modes implying high rates of clonality. Estimates of selfing (S_g_) were overall low with a handful of high values, ranging from 0 to 0.61 (median=0.06). Fifteen populations (28%) were estimated with no selfing. Nineteen populations (36%) showed non-zero estimates under 0.1, 13 populations (25%) between 0.1 and 0.29, and six populations (11%) with estimates between 0.42 and 0.61.

Finally, linkage disequilibria within populations between genotyped SNPs were overall low, with r_d_ values ranging from 0 to 0.51 with a median value of 0.12. Forty-two populations (79%) were under 0.25, and only eight populations (21%) showed linkage disequilibrium between 0.25 and 0.51.

### Analyses of covariations between population genetic indices

The first two components of the principal component analysis on the values of genetic diversity found in the 53 genotyped populations accounted for 70.6% of the total variance between populations (Figure SI4). Non-parametric Kendall partial rank-order correlations between genetic indices (Table SI5) and correlations on the first two principal components from the 17 population genetic indices measured showed three non-collinear groups of associated genetic indices (Figure SI4.A) that are very similar to the theoretical groups of population genetic indices expected to covary with different rates of clonality, selfing and outcrossing in autopolyploids (Stoeckel et al. 2024). A first cluster regrouped G, D*, R, Pareto β and VarF_IS_, indices that are known to be sensitive to clonality (Stoeckel et al. 2021a) but also VarHe, VarHo, and VarAe. This cluster largely explains the first dimension (47.7% of the total variance) of the PCA and was collinear to Σ_CLON_ (also see Figure SI4.B). As expected under partial clonality, VarF_IS_ was negatively correlated to genotypic diversity indices (R and β; Figure SI5). The second cluster regrouped P_ID-SIB_, P_ID-u_, H_E_, A_E_ and H_O_, indices that are linked to the general genetic diversity of populations (Figure SI4.A). The third cluster regrouped S_g_, SE.Sg, MF_IS_ and r_D_, indices that are usually used to identify, rank and estimate rates of selfing versus outcrossing in sexual populations (Castric et al. 2002, Bürkly et al. 2017). The second dimension of the PCA (22.9% of the total variance) was mostly correlated to Ho, Sg, He and r_D_ which was collinear to Σ_SELF_ (Figure SI4.C). The clusters of genetic indices were corroborated by the correlation between Σ_CLON_ and genetic indices (Figure SI5) and Σ_SELF_ and genetic indices (Figure SI6). The correlation across populations between Σ_CLON_ and Σ_SELF_ was negative and highly significant (r_s_ = −0.66; p < 0.001; Figure 2). This correlation within LSI populations was negative and highly significant (r_s_ = −0.65; p < 0.001) while it was no significant in SC populations (r_s_ = −0.32; p =0.289).

**Figure 2:**
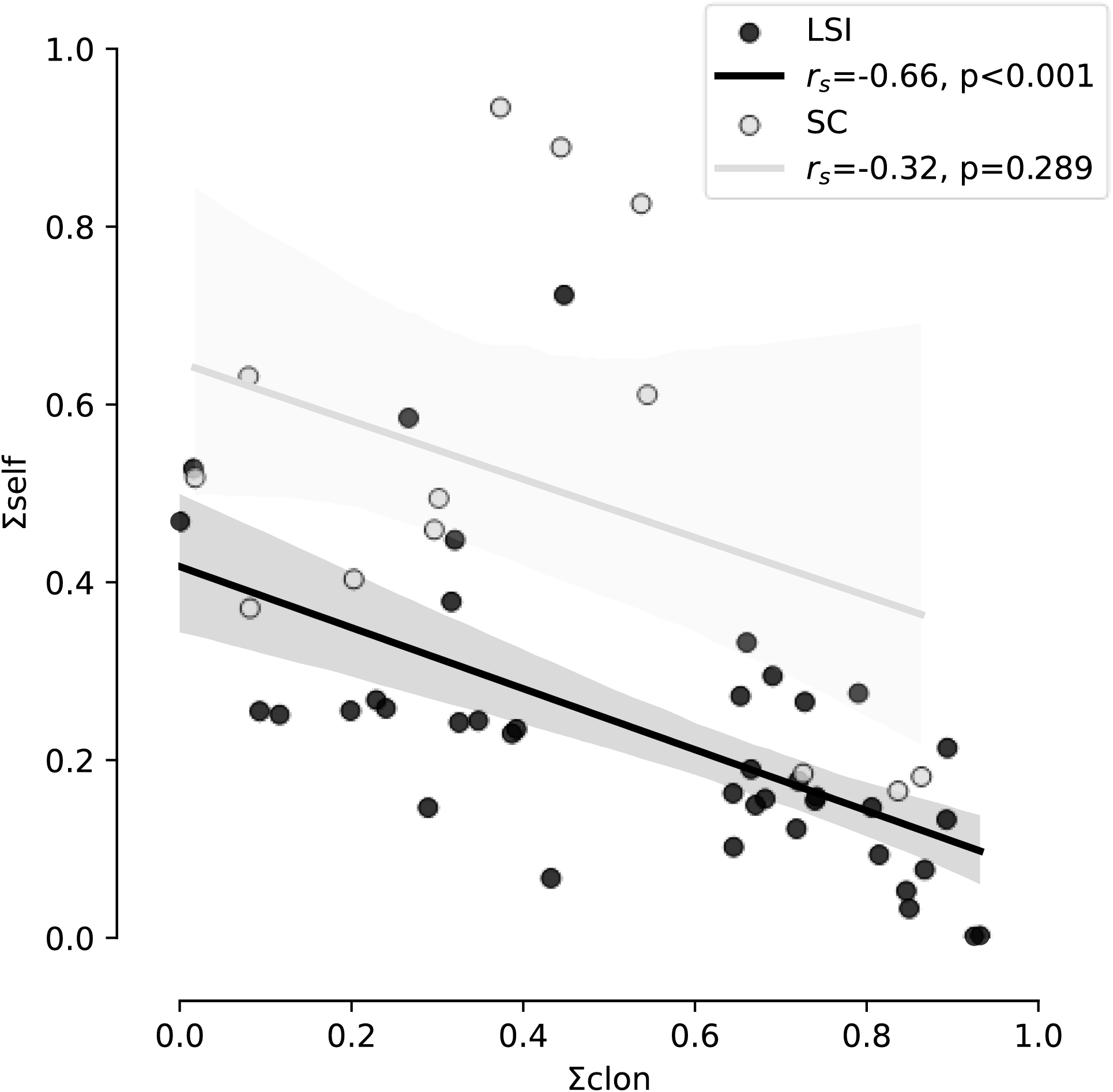
Correlations between Σ_CLON_ and Σ_SELF_ found in 13 SC populations (light grey points and regression line) and 40 LSI populations (black points and regression line). 95% confidence intervals are given for LSI and SC populations in dark grey and light grey, respectively. Spearman rank-order correlation coefficients (r_s_) and probabilities that Σ_CLON_ and Σ_SELF_ would not be correlated (p) are reported for LSI and SC populations.

### Differences in genetic diversity and structure between L- and S-morph populations

The number of ramets per genet was similar in L-morph (LSI) and S-morph (SC) populations (H=1.85, p=0.173, Figure 1 & SI7), as were their genotypic richness (R, H=1.23, p=0.267) and evenness (Pareto’s β, H=1.60, p=0.206; Table SI2). The distributions of other population genetic indices related to clonality were also not significantly different between in L-morph (LSI) and S-morph (SC) populations (VarF_IS_: H=2.94, p=0.086; P_ID-SIB_: H=1.98, p=0.160 and Σ_CLON_ H=2.28, p=0.131; Figure 3). Mean observed heterozygosity (Ho, H=5.27, p=0.022) and their variances (VarHo, H=13.85, p<0.001), mean effective number of alleles per locus (Ae, H=5.13, p=0.024) and its variance over locus (VarAe, H=9.49, p=0.002), MF_IS_ (H=11.85, p<0.001), linkage disequilibrium (r_D_, H=5.55, p=0.018), estimate of selfing rates (Sg, H=12.39, p<0.001) and its standard error over loci (SESg, H=9.58, p=0.002) significantly differed between in L-morph (LSI) and S-morph (SC) populations. S-morph (SC) populations, compared to L-morph (LSI), showed lower mean observed heterozygosity (medians, L-morph: 0.262, S-morph: 0.210) and less variance between loci (medians, L-morph: 2.400, S-morph: 1.859), lower effective number of alleles (medians, L-morph: 1.341, S-morph: 1.283) and less variance between loci (medians, L-morph: 4.419, S-morph: 3.580), higher linkage disequilibrium (medians, L-morph: 0.113, S-morph: 0.152), higher estimates of selfing rate (medians, L-morph: 0.045, S-morph: 0.175) even if with higher standard error between loci (medians, L-morph: 0.058, S-morph: 0.156) and higher Σ_SELF_ (medians, L-morph: 0.222, S-morph: 0.495).

**Figure 3:**
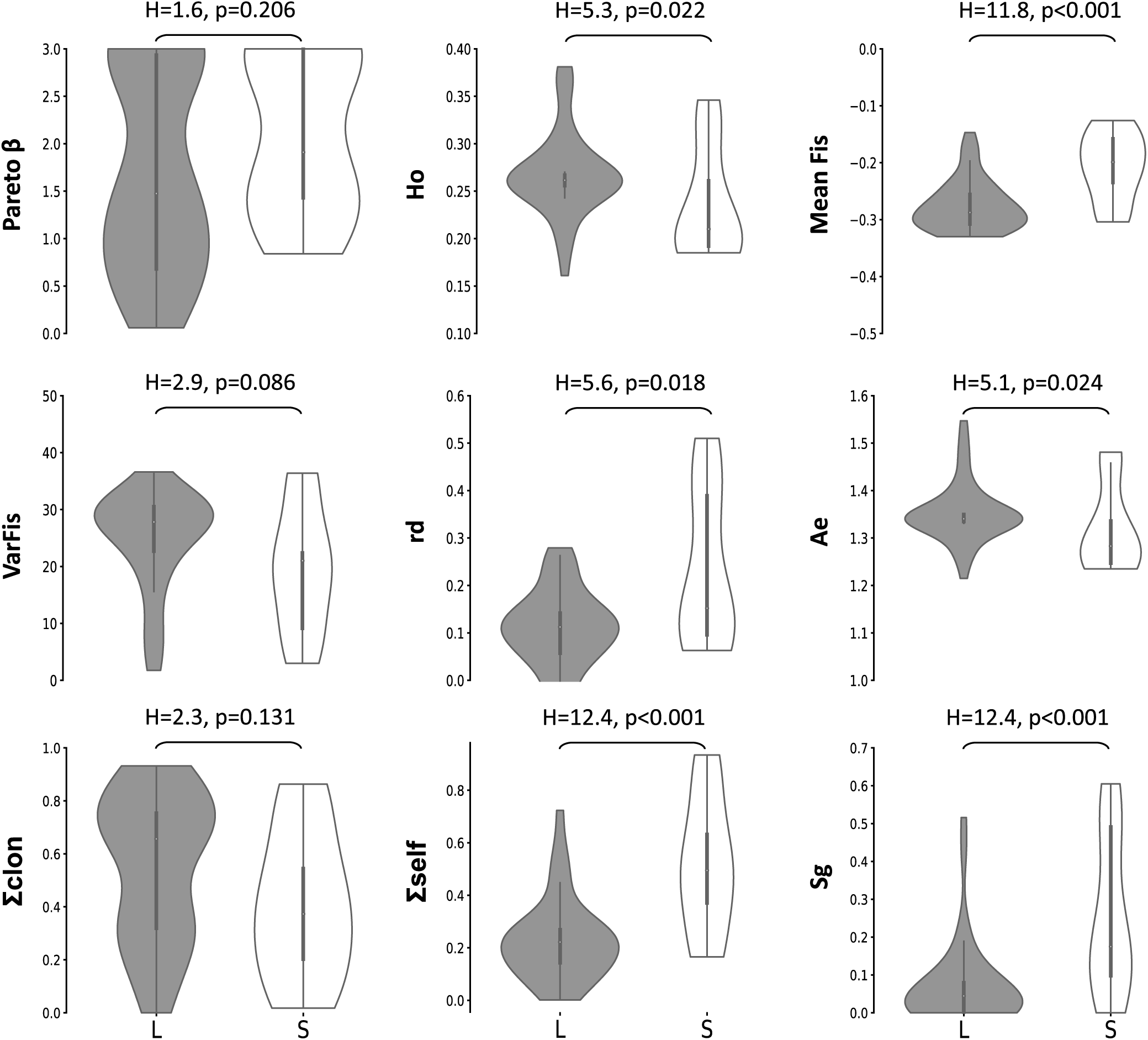
Violin plots comparing distributions of genetic indices in LSI (L, grey) versus SC (S, white) populations. Violin plots are cut for their minimum and maximum values. Kruskall-Wallis tests are reported as H-statistics as well as the probability that LSI and SC population would present the same distribution of genetic indices.

Genetic differentiation (ρ_ST_) among pairs of S-morph (SC) populations (median value of ρ_ST_=0.27) was comparable to the genetic differentiation found among pairs of L-morph (LSI) populations (median value of ρ_ST_=0.26; H=319.37, post-hoc p=0.322; Figure 4). The median of ρ_ST_ between pairs of L-morph (LSI) and S-morph (SC) populations reached 0.65. The distribution of inter-morph ρ_ST_ values differed both from the distribution of pairs of intra-L-morph (LSI) ρ_ST_ values (post-hoc p<0.001) and from the distribution of pairs of intra-S-morph (SC) ρ_ST_ values (post-hoc p<0.001). The minimum spanning tree of genetic distances between individuals computed with *GenAPoPop* showed quite clustered distributions of L-morph (LSI) and S-morph (SC) individuals but also with clear evidence of admixtures between their lineages (Figure 5).

**Figure 4:**
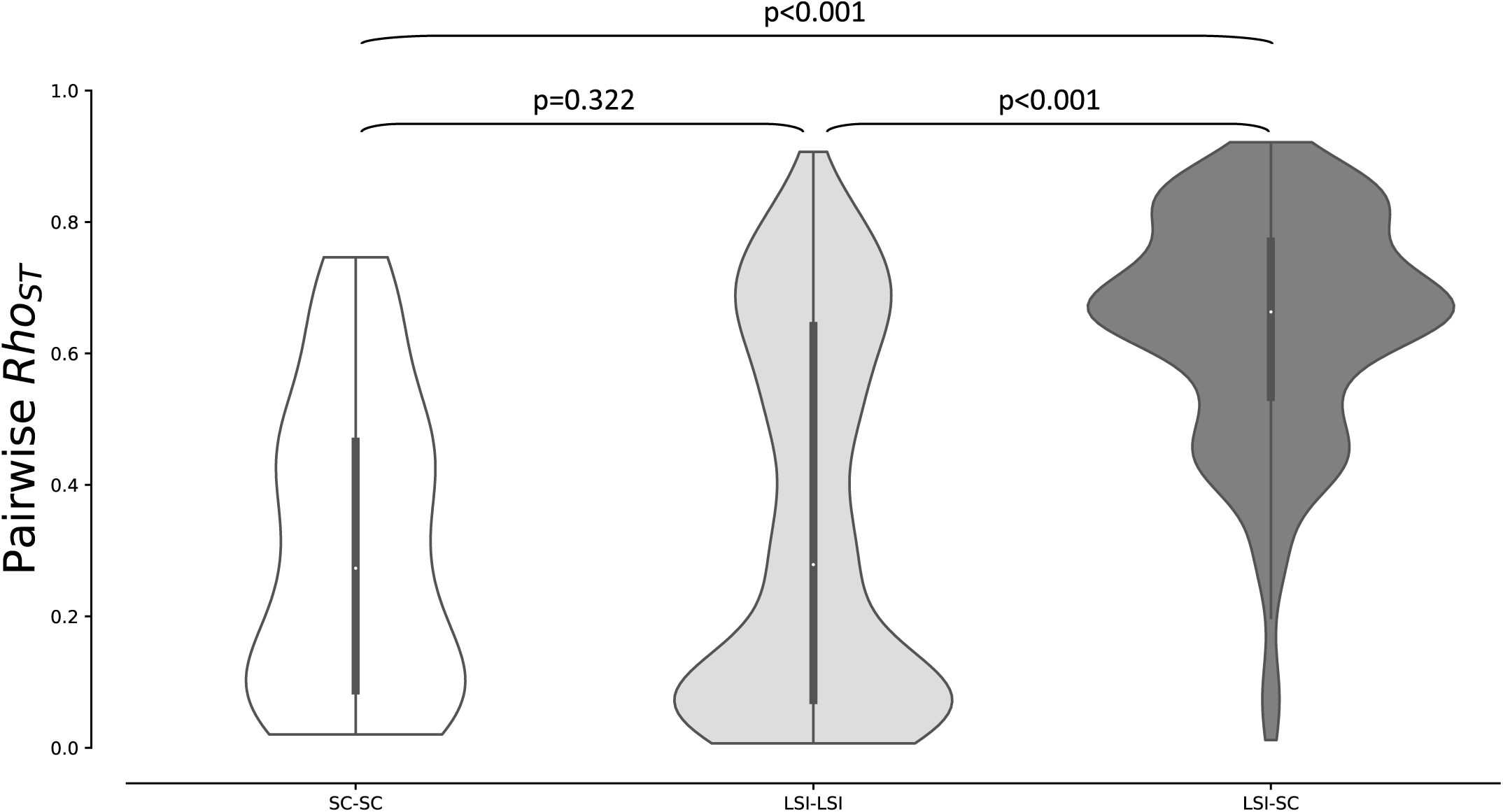
Distributions of pairwise rho_st_ within pairs of SC populations (white violin plot), pairs of LSI populations (light grey violin plot) and between pairs of LSI and SC populations (dark grey violin plot). Probabilities that pair of non-parametric distributions are equal are reported.

**Figure 5:**
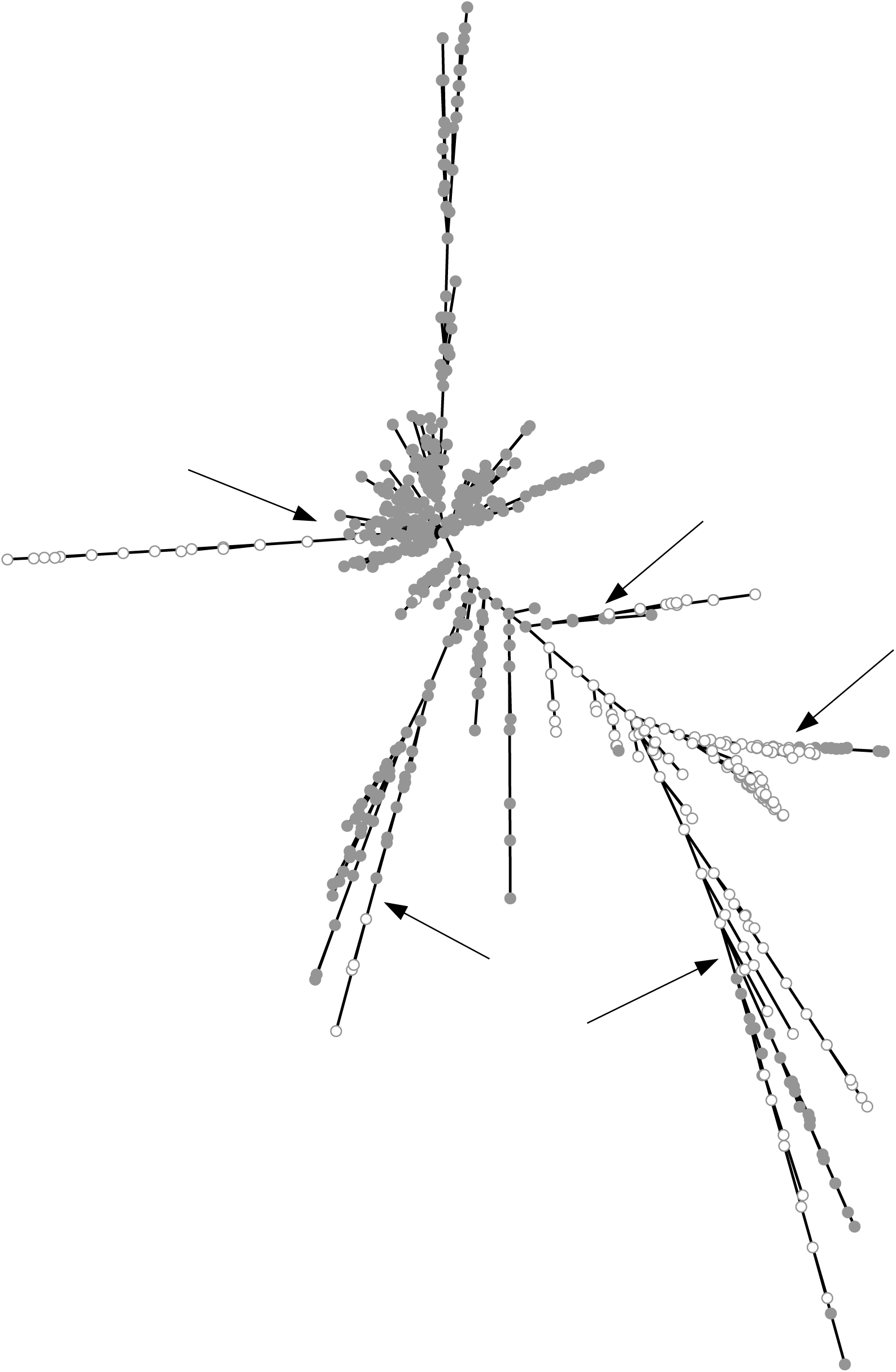
Minimum spanning tree of the genetic distances (number of different alleles) between L-morph (LSI, grey points) and S-morph (SC, white points) individuals. Arrows indicate some of the evidences of admixture between LSI and SC lineages.

## Discussion

We used population genetics on autotetraploid SNPs with allele dosage to assess the rate of clonality, of selfing and of outcrossing (i.e., reproductive mode) of 53 populations of *Ludwigia hexapetala* subsp. *grandiflora* (*Lgh*) recently colonizing western European watersheds.

To achieve this goal, we developed reproducible codominant molecular markers (SNPs) that enabled allele dosage-based genotyping of sampled individuals. We then used tailored computational analyses to perform confident population genetic analyses in autopolyploids. These two methodological steps allowed solving the remaining challenges to perform population genetic analyses in autopolyploids (Dufresne et al. 2014), including *Lgh*. The resulting framework can be applied to any autopolyploid species.

*Lgh* develops two floral morphs respectively associated with a Late-acting self-incompatible system (L-morph) and a self-compatible system (S-morph, Portillo-Lemus et al. 2021, 2022). Interestingly, the sampled populations in western Europe showed either L- or S-morph resulting in two groups of sampled populations: one group of 40 L-morph (LSI) populations and one group of 13 S-morph (SC) populations. They live in similar ecological conditions (Portillo-Lemus et al. 2021) which thereby provides a rare opportunity to characterize the genetic consequences of an LSI system compared to a group of SC populations in the same species and ecological context.

**F**ive L-morph and two S-morph water primrose populations separated by 50 to 150 km from one to the other in France we karyotyped, all had 80 chromosomes corresponding unequivocally to the species *Ludwigia grandiflora* subsp. *hexapetala* and thus didn’t find any individual with 48 chromosomes corresponding to *Ludwigia grandiflora* subsp. *grandiflora*. These results agree with previous observations of Dandelot et al (2005) and Barloy et al. (2024) in France, Bou Manobens et al. (2019) in northern Spain and Armitage et al. (2013) in Great Britain.

### Most of the genetic variance explained by clonality and selfing among *Lgh* populations

The correlations of population genetics indices on the 53 genotyped *Lgh* populations (Figure SI4) were congruent with previous theoretical predictions on the variations of genetic indices with different rates of clonality and selfing for autopolyploid populations (Stoeckel et al. 2024). Identical covariations were already predicted by Wright-Fisher-like models adapted for diploid (Stoeckel et al. 2021a) and haplodiplontic life-cycles (Stoeckel et al. 2021b), and validated respectively using multiple field populations of marine phanerogams for diploids (Arnaud-Haond et al. 2020) and of brown, green, red algae and mosses for haplodiplontics (Krueger-Hadfield et al. 2021). All these predictions and observations argue for the primary importance of reproductive mode, especially of high clonality, to drive variation in genetic diversity within populations (Duminil et al. 2007). They support the methodological importance of accurate assessment of rates of clonality and of selfing before starting to interpret genetic diversity.

### Dominant clonality within the western European *Lgh* populations

Previous field and lab observations of peripatric *Lgh* populations reported massive production of dispersing vegetative propagules, rapid expansions of patches, and an important capacity for spontaneous cutting-planting (Dandelot et al. 2005, Thouvenot et al. 2013, Grewell et al. 2016, Skaer-Thomason et al. 2018a,b). Okada et al. (2009) genotyped around 800 individuals sampled in 27 *Lgh* populations in Californian wetlands using a set of eight AFLP markers and reported an extremely reduced clonal diversity: 95% of the samples had the same genotype and 18 populations over 27 (67%) supported a single AFLP-genotype. All over the UK, only two haplotypes on 14 sampled stems were found using chloroplast sequences (Armitage et al. 2013). These haplotypes were even shared with some *Lgh* samples invading California. With no measure of the probability of identity to assess the marker set, these results could be due to the lack of resolution of the markers used resulting in an artificially elevated measure of clonality (Waits et al. 2001, Villate et al. 2010). In any case, all these studies concluded that invasive populations of *Lgh* in Europe and in USA reproduce by exclusive clonality, with a very narrow base of ancestral clones or being monoclonal (Dandelot et al. 2005, Okada et al. 2009, Thouvenot et al. 2013, Grewell et al. 2016).

In our study, we chose to develop and use SNPs rather than AFLP markers. The latter can be challenging in terms of precisely sizing fragments, leading to suboptimal reproducibility, particularly across different platforms (Fry et al. 2009). The 36 SNPs we developed are easily reproducible, accurate enough to distinguish between offspring of sibling mating as evidenced by the probabilities of identity we obtained, and less expensive.

With these SNPs, we found typical genetic signatures of high clonality, including clear occurrence of replicated genotypes and mean negative F_IS_ values with high interlocus variances, in all the 53 western European populations we genotyped. Globally, gene diversities measured in these invasive *Lgh* populations were in the higher range of values commonly found for SNPs (Fischer et al. 2017, Schmidt et al. 2021). Such levels of gene diversity are in line with strong rates of clonality that buffer the loss of alleles due to genetic drift (Reichel et al. 2016, Stoeckel et al. 2021b). Among the 58 MLGs with replicates, seven MLGs were found in a single population and 51 MLGs were found in multiple sites. Clonal propagules of *Lgh* are known to disperse carried by river current and by birds (zoochory: seedlings have the ability to stick to feathers; Grewel et al. 2016). Clonal reproduction by rhizomes at the local scale and by the dispersal of clonal propagules would thus be the main source of population growth and invasive spread of *Lgh* in western Europe.

Our results also report a previously-underestimated genotype diversity in the invasive populations in western Europe. We found 462 distinct MLGs (genets) out of 795 individuals within these populations and the large majority of these MLGs (404, thus 88%) were only sampled once in the 53 populations. Our results are still congruent with Dandelot (2004) unpublished measures obtained in three Mediterranean populations using inter-simple sequence repeats (ISSRs). This data reports three and seven genotypes over 11 sampling units within two *Lgh* populations and two genotypes over nine sampling units in a third population, all sampled in the south-east of France. These results could reveal contrasted situations between *Lgh* populations introduced in the USA and continental Europe.

### Between 10 and 40% of sexual reproduction in *Lgh* populations in western Europe

Beyond the qualitative indication of dominant clonality, we then aimed at estimating the rates of sexuality in each of these 53 populations. All showed small to medium values of linkage disequilibrium between SNPs. Such values are expected in highly but non-exclusive clonal populations with large population sizes (Navascues et al. 2010, Stoeckel et al. 2021a). Thirty-two populations showed Pareto β values of their distributions of clonal sizes under 2, which are only found in theoretical populations with rates of clonality higher than 0.8 (Stoeckel et al. 2024). All 53 populations showed mean negative F_IS_ with high interlocus variance, with values observed in theoretical populations with rates of clonality higher than 0.6 but under 0.9 (Balloux et al. 2003, Stoeckel & Masson 2014, Reichel et al. 2016, Stoeckel et al. 2024). All these values of population genetic indices were consistent with the interpretation that *Lgh* populations in western Europe must reproduce with effective rates of clonality between 60% and 90%, thus with 10% to 40% sexuality (De Meeus et al. 2006, Arnaud-Haond et al. 2020, Stoeckel et al. 2024). These estimates were also supported by the local evidence of recombination between clonal lineages, and even between L- morph (LSI) and S-morph (SC) lineages at the scale of a watershed (Figure 5). The diversity of genotypes detected in western Europe is thus indicative of rare but significant local sexual events rather than of a large clonal diversity that would have maintain and propagate by exclusive clonality since its introduction. Our results newly advocate that sexual reproduction should not, therefore, be overlooked in these invasive populations, especially in management plans.

### Genetic consequences of LSI compared to self-compatible populations

The late-acting SI system remains one of the less studied breeding systems among the mechanisms favouring outcrossing in plants (Gibbs 2014). Its efficiency to favour outcrossing and its consequences on genetic diversity within populations, especially considering its low but common failures, were not yet deciphered and not yet compared to SC populations in the same ecological conditions, as already previously explored for gametophytic and sporophyte SI systems (Busch 2005, Koelling et al. 2011).

The maintenance of SI systems is one of the most intriguing evolutionary puzzles (Porcher & Lande 2005). Indeed, SI systems are fated to breakdown because SC individuals present the advantage of reproductive assurance when compatible partners are limited, especially in peripatric conditions (Eckert et al. 2006). This advantage is even absolute when no compatible partners are available within pollination range, as we commonly found in *Lgh* populations in western Europe (Figure SI1). Conversely, outcrossing imposed by SI systems decreases the probability of expressing deleterious mutations in descendants as compared to selfing with the same genetic background (Rice 2004, Navascues et al. 2010).

We found that the 40 populations with L-morph individuals (LSI) had a higher number of effective alleles, higher gene diversity, higher observed heterozygosity and less linkage disequilibrium than the 13 populations with S-morph individuals (SC; Figure 3).

These genetic differences are very unlikely to directly result from the consequences of the LSI system hitchhiked to the whole genome in L-morph individuals and populations. Indeed, it would imply either that all the 36 SNPs would be physically linked to the genes under negative frequency- dependent selection coding for the LSI or that *Lgh* outcrossed for many generations in small population sizes (Glémin et al. 2001, Navascues et al. 2010). *Lgh* in western Europe develops including far more than thousands of stems per local population (Portillo-Lemus et al. 2021) and the linkage disequilibrium values also argued for large effective population sizes. We found similar estimated rates of sexuality and of clonality between L-morph and S-morph populations, using Σ_CLON_ and all indices sensible to clonality (R, Pareto β and VarFis; Figure 3). We however found significant difference in estimates of selfing between L-morph (LSI) and S-morph (SC) populations, and in Σ_SELF_ and in all indices sensible to selfing (Ae, linkage disequilibrium, Ho, mean Fis) revealing typical signatures of higher selfing rates in S-morph populations. Hardy (2016) method estimated a median selfing rate (Sg) of 0.18 in S-morph populations versus 0.05 in L-morph populations. Consequently, the genetic differences we found between S-morph and L-morph populations may thus rather be due to the effects of selfing impacting S-morph (SC) populations than due to outcrossing protecting the loss of genetic diversity or rates of clonality in L-morph (LSI) populations.

### Advantages of dominant clonality with preferential allogamy in invasive *Lgh* populations

Uniparental clonal reproduction (including clonality and selfing) may help the demographical maintenance of plants spreading out of their native range with limited or even without compatible or less related sexual partner at pollination distance (Baker’s conjecture: Barrett et al. 2008, Pannell et al. 2015). If they rather reproduce using selfing, their descendants increase the probability to express inbreeding depression and to lose heterozygosity. Some invasive populations develop with low genetic diversity (He et al. 2024), questioning on the biological and environmental factors that may explain their success (i.e., the genetic paradox of invasions, Allendorf & Lundquist 2003). But many other plants spread out of their native ranges with substantial genetic diversity and using reproductive modes that favour outcrossing (Roman & Darling 2007, Forsman 2014), like *Lgh* populations in western Europe, mostly when developing in harsh and stressful conditions (Fox & Reed, 2011) or when the costs of inbreeding depression expressed by selfing are superior to the benefits of reproductive assurance (Layman et al. 2017).

Peripatric populations of *Lgh* in western Europe seem to solve all these problems and paradox by mixing clonality with preferential allogamy: clonality allows the local maintenance and spreading of population without losing heterozygosity and genetic diversity, and subtle but significant sexuality with preferential allogamy, favoured by LSI and faster growth of crossed-pollen tubes, enables recombination between lineages favouring the emergence of locally adapted genomes with potential higher vigour and fertility (heterosis: Darwin 1876, Lippman & Zamir 2007, Birchler et al. 2010).

This reproductive mode, i.e. dominant clonality with preferential allogamy, is common in plants (Vallejo-Marin et al. 2010). The micro-physiological mechanism(s) slowing down the growth of self-pollen tubes, rather than blocking them, in simultaneous monoecious and hermaphrodite SC species and resulting to favour allogamy when compatible pollen is available may also be common in plant, although potentially overlooked (Glover 2007, Nasrallah 2017). In such peripatric conditions, selfing would thus only present the limited or transient advantage to produce seeds that can maintain in local seed banks and with different dispersal properties compared to clonal propagules. The limited benefit of selfing in this species may explain why, based on photos collected on the web, L-morph (LSI) individuals seem more frequent in peripatric than in native populations, and why in western Europe, around 76% of the populations are L-morph (LSI) against 24% of S-morph (SC; Portillo et al. 2021). Our results thus call for measuring the true proportions of LSI and SC in invasive versus native worldwide *Lgh* populations with estimation of their rates of clonality and of selfing.

### Unusual selfing syndrome in *Lgh* populations

Highly selfed populations of different plant species tend to share similar morphology and functions resulting in a set of traits called *selfing syndrome*, including reduced flower size (Darwin 1876, Tsuchimatsu & Fujii 2022). Selfing syndrome, including reduced flower size, seems to evolve rapidly, as observed for example in five generations in *Mimulus guttatus* (Bodbyl-Roels & Kelly 2011), in four generations in *Silene latifolia* (Delph et al. 2004), in three generations in *Phlox drummondii* (Lendvai & Levin 2003) and after only two generations in *Eichhornia paniculata* (Worley & Barrett 2000).

On the contrary, S-morph (SC) individuals in *Lgh* develop larger flowers than allogamous L-morphs (Portillo-Lemus et al. 2021). However, S-morph (SC) individuals and populations dominantly reproduced using clonality and produced selfed offspring only when lacking crossed pollen. Both clonality and faster growth of crossed-pollen tubes may delay or even compromise the emergence of the first steps of a selfing syndrome. These two traits may call for further studies on the emergence of selfing syndrome in partially clonal and selfed populations.

## Conclusion

We found that peripatric populations of *Lgh* in western Europe reproduced using dominant clonality with limited but stable significant sexuality with preferential outcrossing, in nearly all populations, within a large clonal diversity. *Lgh* is one of the most invasive aquatic plants in the world, and considerable efforts are made to limit its deleterious effects in the newly colonized ecosystems (Thouvenot et al. 2013, Grewell et al. 2016, Portillo-Lemus et al. 2021). The rare sexual events, allogamous when possible, occurring in invasive peripatric populations of *Lgh* may favour the emergence of new genotypes, more adapted to local conditions. Managers should thus chiefly concentrate their actions on the most sexual populations, and on the contact zones between S- and L-morph populations in France. We also found variations of the rates of clonality and of selfing among *Lgh* populations. Considering the importance of reproductive modes on the dynamics and evolution of populations (Duminil et al. 2007, Ellegren& Galtier 2016, Glémin et al. 2019), our results advocate that management actions should consider the local effective reproductive modes to control *Lgh* population by population. Finally, knowing the reproductive modes of populations, the distribution of clones and the self-compatibility across western European populations now allow deciphering and interpreting their population structure, identifying their origin and routes, and predicting their possible short-term dynamics.

## Data Accessibility Statement

The data that support the findings of this study are openly available on Zenodo (https://doi.org/10.5281/zenodo.12760022).

## Benefits Generated

Benefits from this research accrue from the sharing of our data and results on public databases as described above.

## Acknowledgements

We thank Michel Bozec (UMR DECOD), Diane Corbin (FRAPNA Loire-Ecopôle du Forez), Guillaume Le Roux (Reserve Naturel Val d’Alier Châtel-de-Neuvre), Fabrice Dejoux (Service Agriculture Environnement Roannais Agglomération and Jordi Bou Manobens (University of Girona, Spain) for helping to collect *Lgh* samples. We thank Olivier Coriton and Virginie Huteau for their help acquiring karyotypes.

We warmly thank Ingrid Leveque, Marie-Therese Delaroche, Gervaise Fevrier and Patricia Nadan for their administrative support and facilitation.

## Funding information

This work was supported by FEDER Région Centre-Val de Loire, by Agence de l’eau Loire-Bretagne (grant Nature 2045, programme 9025, AP 2015 9025) and by the French National Research Agency (project Clonix2D ANR-18-CE32-0001). FEDER funded the doctoral grant of L. Portillo-Lemus and salary of Marilyne Harang. Postdoctoral grant of Ronan Becheler was funded by Clonix2D ANR-18-CE32-0001.

## Conflict of interest

The authors declare that they have no financial conflicts of interest based on the content of this article.

## Author contributions

DB and SS laid the foundation of this work, conceived the study and were responsible for funding applications. LPL, MH and DB collected samples and performed field observations. LPL and DB read karyotypes and counted chromosomes. EP and DB conceived and supervised RAD4SNP and Hi-Plex methodology, BJP and DJP contributed to define the SNP set. LPL, MH and ALB performed RAD4SNP approach to develop the SNPs. LPL and MH extracted DNA and genotyped the samples, SMC and RCV sequenced the data. GL performed the bioinformatic analyses and SS computed the Bayesian allele dosages. DB, LPL and SS performed early data explorations at the end of LPL PhD thesis that was supervised by DB and SS. RB and SS performed computations and analyses, tracked and managed the bibliography, produced tables, figures and interpretation. DB, RB and SS wrote the manuscript. All authors read and approved the final manuscript.

## Supplementary Information (SI)

**Supplementary information1:**
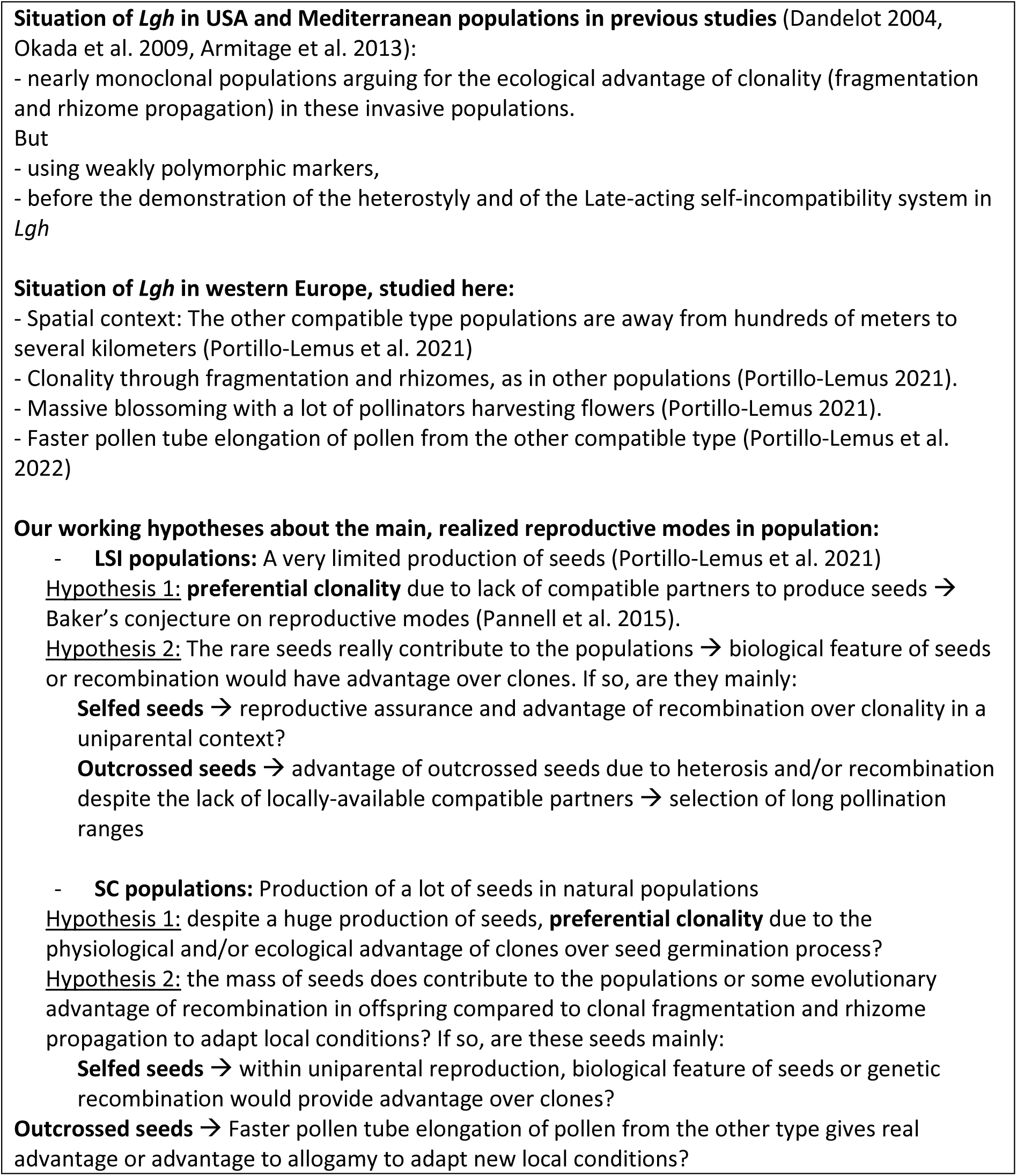
Synthesis of the questionings of our study to estimate reproductive mode of 53 populations of invasive Lgh using genetic diversity on the autotetraploid part of its genome.

**Table SI1:** Hypotheses about the influence of clonality and selfing on the expected variations of genetic indices.

| Genotypic or genetic index | Range of values | Main driver | Behavior | references |
| --- | --- | --- | --- | --- |
| $\beta^*$ | $\in [0; +\infty]$ | Rate of clonality | ↘ when $c \nearrow$<br>$\beta \rightarrow 0$ when $c \rightarrow 1$ | Halkett et al. 2005, Arnaud-Haond et al. 2007, 2020, Stoeckel et al. 2021 |
| $\text{varF}_{IS}$ | $\in [0; +\infty]$ | Rate of clonality | ↗ when $c \nearrow$<br>$\text{VarF}_{IS} \rightarrow 0$ when $c \rightarrow 0$ | Stoeckel & Masson 2014, Stoeckel et al. 2021 |
| $r_d$ | $\in [0; 1]$ | Rate of selfing | slightly ↗ when $c \nearrow$<br>↗ when $s \nearrow$<br>$r_d \rightarrow 1$ when $s \rightarrow 1$ | Weir and Cockerham 1973, Navascues et al. 2010, Halkett et al. 2005 |
| $H_o$ | $\in [0; 1]$ | Rate of selfing | ↘ when $s \nearrow$<br>$H_o \rightarrow 0$ when $s \rightarrow 1$ | David et al. 2007, Halkett et al. 2005 |
| $F_{IS}$ | $\in [-1; 1]$ | Both rates | ↘ when $c \nearrow$<br>$F_{IS} \rightarrow -1$ when $c \rightarrow 1$<br>$F_{IS} \rightarrow +1$ when $s \rightarrow 1$ | Stoeckel & Masson 2014, Balloux et al. 2003, Stoeckel et al. 2021 |
| $H_E$ | $\in [0; 1]$ | Rate of selfing | ↘ when $s \nearrow$<br>$H_E \rightarrow 0$ when $s \rightarrow 1$ | Glémin et al. 2001, Ritland & Ganders 1987 |
| $P_{ID-SIB}$ | $\in [0; +\infty]$ | Both rates | ↘ when $c \nearrow$<br>↘ when $s \nearrow$ | Waits et al. 2001 |
| $S_g^{**}$ | $\in [0; 1]$ | Rate of selfing | ↗ when $s \nearrow$ | David et al. 2007, Hardy et al. 2016 |

\* similar behavior of the clonal richness R
\*\* inferred values of s, according to the method of Hardy (2016)

**Table SI2:**
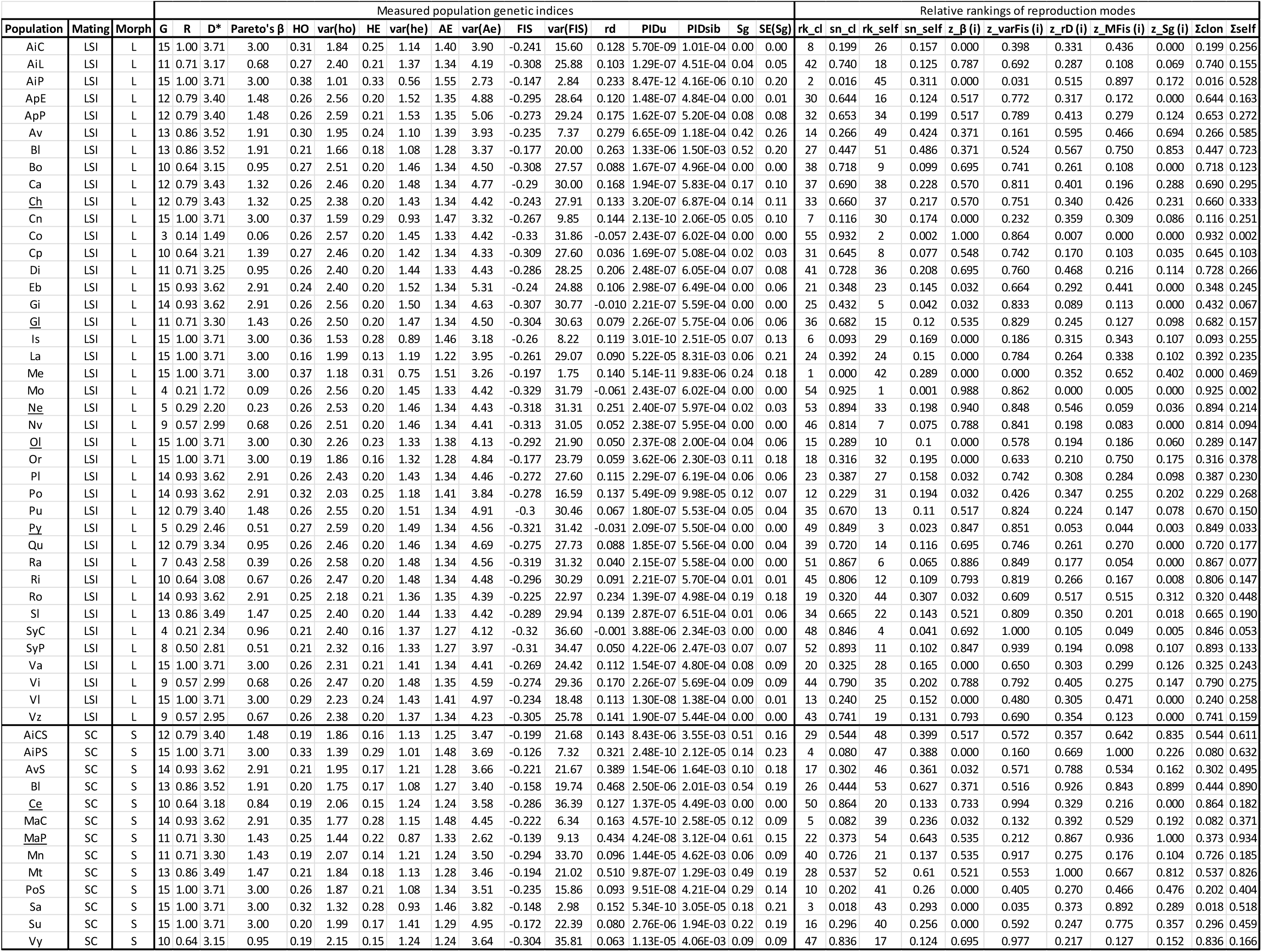
All genetic and genotypic indices computed on the 53 sampled populations. Each index was obtained from 15 individuals sampled in each population. (See spreadsheet supplementary material). *Mating* column indicates if the population was supposed to expressed a late-acting self-incompatible system (LSI) or be self-compatible (SC), the seven underlined populations being directly tested by controlled crosses using 15 plants randomly sampled in the population. *Morph* indicates the floral morph of the population. *G* is the number of multi-locus genotypes genotyped on the 15 sampled individuals per population; *R* is the genotypic diversity; *D\** clonal heterogeneity; *Pareto’s β* the clonal evenness; *HO*, the mean observed heterozygosity; *var(ho)*, the variance of the observed heterozygosity among loci; *HE*, the mean expected heterozygosity; *var(he)*, the variance of the expected heterozygosity among loci; *AE*, the mean effective number of alleles among loci; *var(Ae)*, the variance of the effective number of alleles among loci; *FIS*, the mean inbreeding coefficient; *var(FIS)*, the variance of the inbreeding coefficient among loci; *rd*, the population index of multilocus linkage disequilibrium; *PIDu* and *PIDsib*, unbiased probability of identity under panmixia and between sibs respectively; *Sg*, the estimate of selfing and *SE(Sg)* the standard error of estimate of selfing. Then, the relative rankings of populations according to different indices and the relative rankings of reproduction modes (see Material and Methods). Full table is available at https://doi.org/10.5281/zenodo.12760022, tabulation “*Genetic indices per pop*”.

**Table SI3:** Summary of the allele dosage. First subtable lists populations with individual genotyped with at least one individual with missing genotype at one locus. Second subtable reports the frequency of single SNP genotypes with posterior probability of allele dosage strictly superior to 70%. Third subtable reports the frequency of individuals with their 36 SNP genotypes assigned with posterior probability superior to 70%.

| Population with missing genotype | nb_missing_genotypes | nb_individuals_concerned |
| --- | --- | --- |
| Bléré_Ls | 1 | 1 |
| Chambéon_écopôle_du_Forez | 2 | 2 |
| Le_Port_Ls | 1 | 1 |
| Plougonver_Léguer | 1 | 1 |
| Quevillon | 1 | 1 |
| Rennes_apigné_prairie | 1 | 1 |
| Roost-Warendin | 1 | 1 |
| Saint_Aignan_Couflons_Ls | 1 | 1 |
| Vichy | 1 | 1 |

| Over single SNP genotypes | N | %age |
| --- | --- | --- |
| single SNP genotype P(cad)>70% | 28610 | 0.99965 |
| single SNP genotype P(cad)≤70% | 10 | 0.00035 |
| total | 28620 | 1 |

| Over individuals | N | %age |
| --- | --- | --- |
| nb_individuals_with_full_genotype | 785 | 0.98742 |
| nb_individuals_with_one_missing_genotype | 10 | 0.01258 |
| total | 795 | 1 |

**Table SI4:** Distribution of the different repeated Multi-Locus Genotypes (MLGs) found with 36 SNPs among populations considering their mating system (LSI or SC). Figure 1 presents a summarized plot of this table. (See spreadsheet supplementary material).

**Table SI5:**
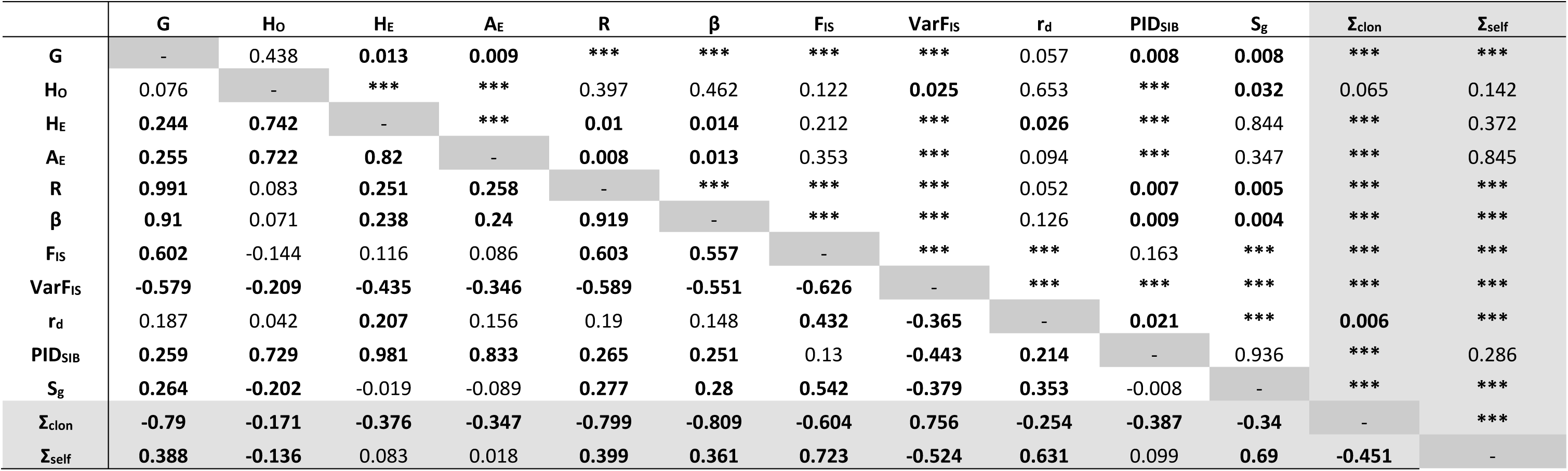
Correlations between genetic indices using non-parametric Kendall partial rank-order correlation. Below the diagonal, the values of the coefficient of correlation τ are consigned. Above, the related p-values are provided. Values inferior to the threshold 0.05, and the related coefficient τ were bold. The parameters Σ_CLON_ and Σ_SELF_ correspond to the sum of normalized indices sensitive to clonality and selfing, respectively.

**Figure SI1:**
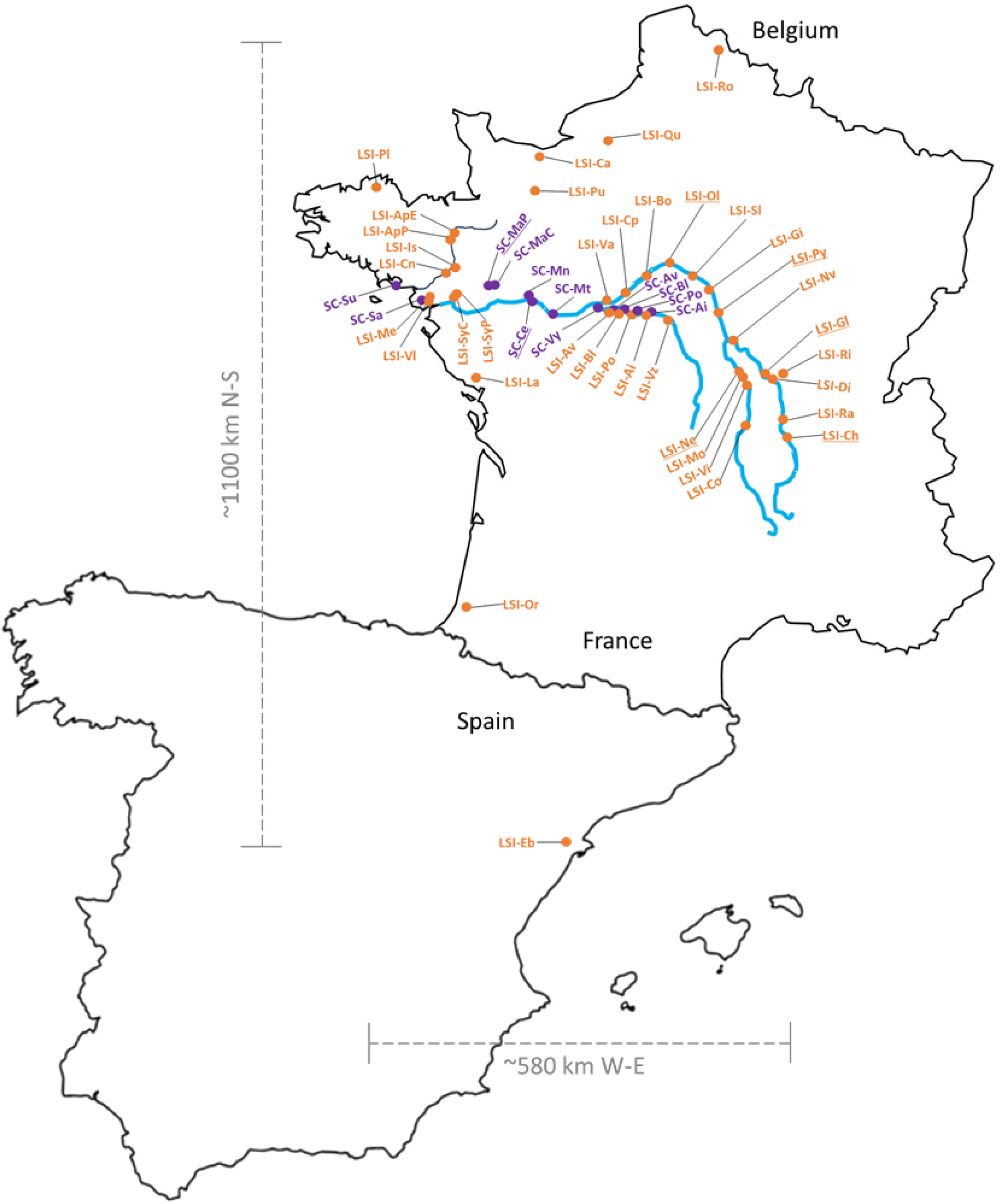
Map of the locations of the 53 sampled populations across France and Spain. In orange, populations with only L-morph individuals, supposed to mate under the control of a late-acting self-incompatible (LSI); In purple, populations with only S-morph individuals, supposed to be self-compatible (SC). In light blue, the Loire river system. In grey, the North-to-South (N-S) and West-to-East (W-E) scale distances of the sampled populations. Underlined, the seven populations in which 15 individuals were tested for the self-incompatibility (Portillo-Lemus et al. 2021) and one individual was karyotyped to count the chromosome numbers.

**Figure SI2:**
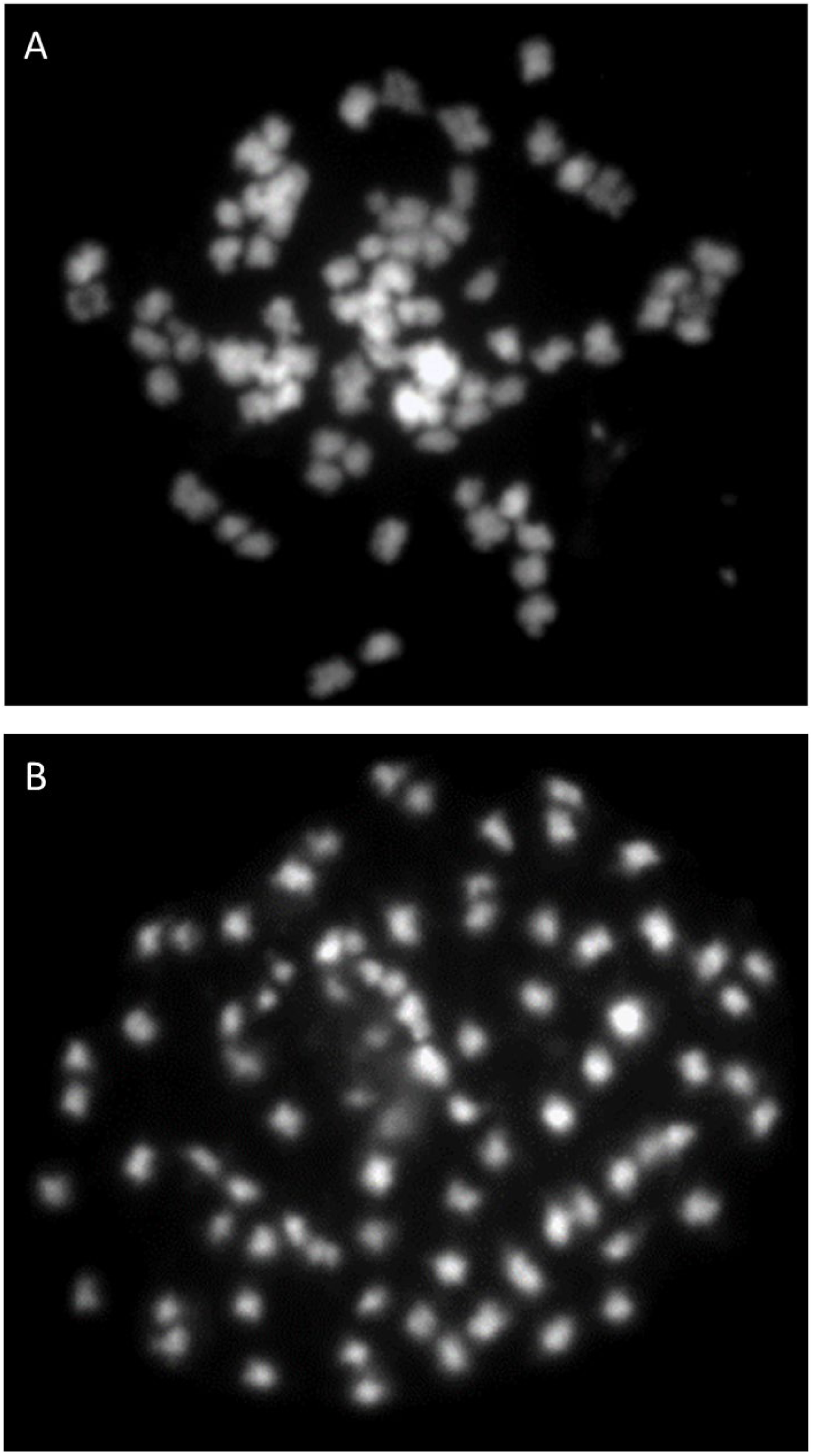
Photos of somatic metaphase chromosomes indicating that L- and S-morph individuals that succeed to mate and to give full fruit set and 100% viable first- and second-generation descendants (Portillo-Lemus et al. 2021) also present the same number of chromosomes (2n=10X=80), belonging to the same taxon *Ludwigia grandiflora* subsp. *hexapetala*. We still didn’t find *Ludwigia grandiflora* subsp. *grandiflora* karyotype in France (2n=6X=48, see Dandelot 2005, Barloy et al. 2024). **A**: somatic metaphase chromosomes from L-morph (2n=10X=80); **B**: somatic metaphase chromosomes from S-morph (2n=10X=80).

**Figure SI3:**
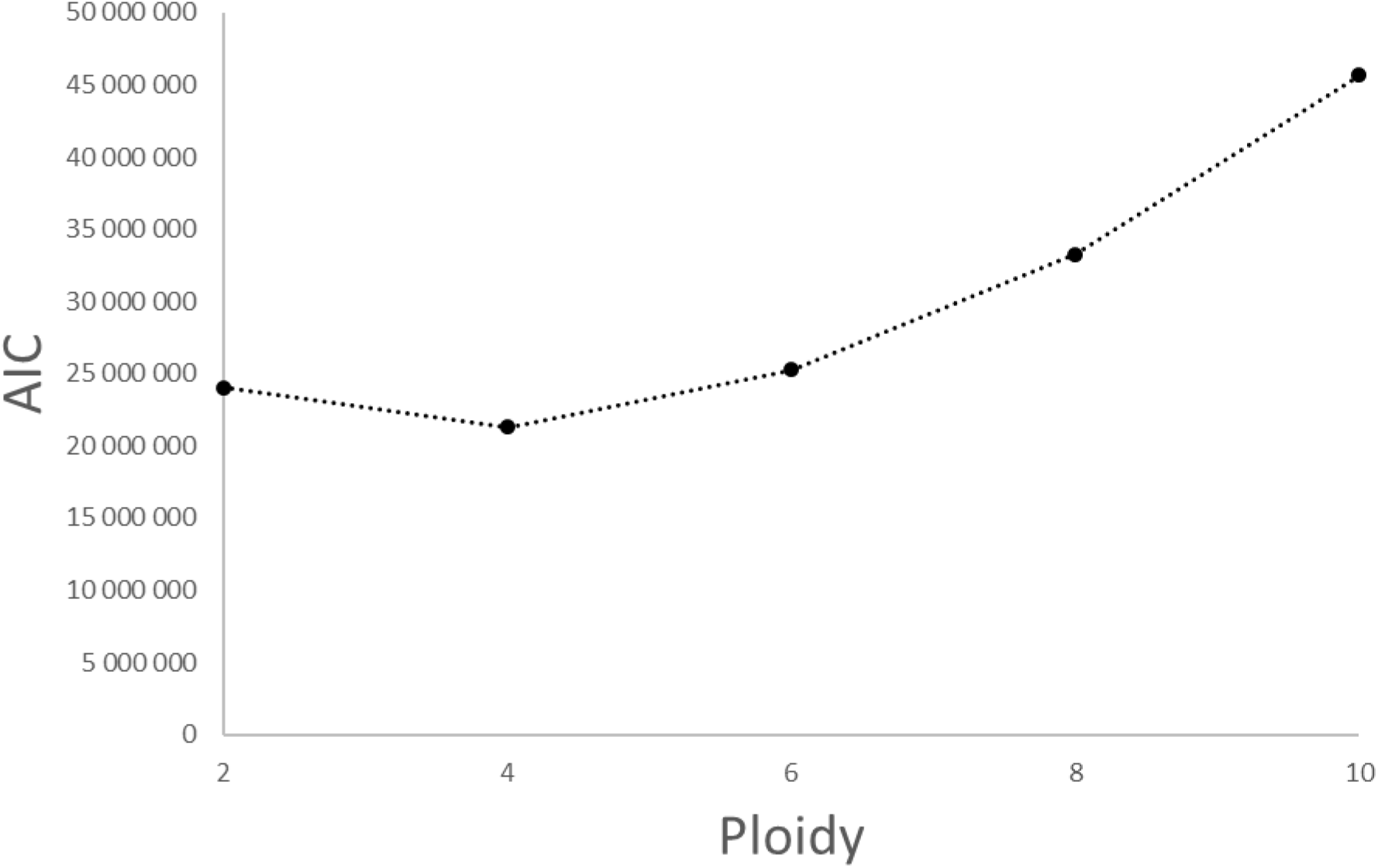
Akaike’s information criterion (AIC) as function of the ploidy level on the likelihoods of the sequenced allele countings over all individuals. Tetraploidy, a ploidy 4x, presents the lowest AIC, and thus the best supported model to explain distribution of sequenced allele countings over all individuals, which confirm our initial SNP development to be sure of their ploidy level.

**Figure SI4.A:**
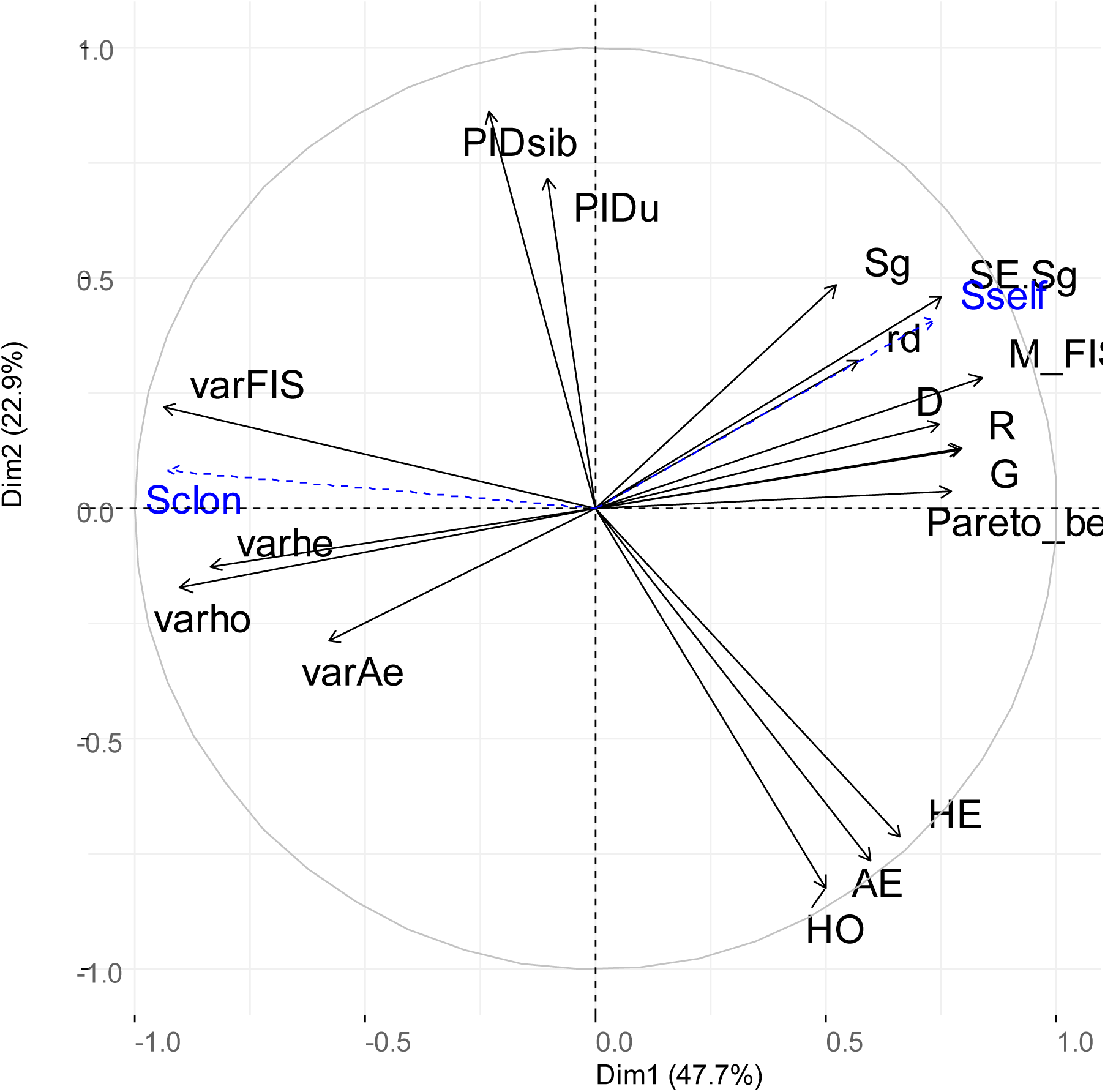
Correlation circles of the principal component analysis on the 17 centered and scaled genetic indices measured in 53 *Lgh* populations in western Europe. The first horizontal dimension accounting for 47.7% of the variance regroups indices related to clonality (number of genotypes *G*, genotypic diversity *R*, clonal heterogeneity *D*, clonal evenness *Beta_Pareto*, variance of Fis *varFis*, variance of expected heterozygosity *varHe*, variance of observed heterozygosity *varHo* and variance of effective number of alleles varAe between loci). The second vertical dimension accounting for 22.9% of the variance with a main contribution of estimate of selfing Sg. Other indices, *i.e.*, mean observed heterozygosity *Ho*, mean expected heterozygosity *He*, effective number of alleles *Ae*, inbreeding coefficient *Mfis*, linkage disequilibrium *rd*, standard error of estimate of selfing *SE.Sg*, unbiased probability of identity under panmixia *PIDu* and between sibs *PIDsib*. In blue and dashed lines, the predictions of Σclon (*Sclon*) and Σself (*Sself*) provided as supplementary variables.

**Figure SI4.B:**
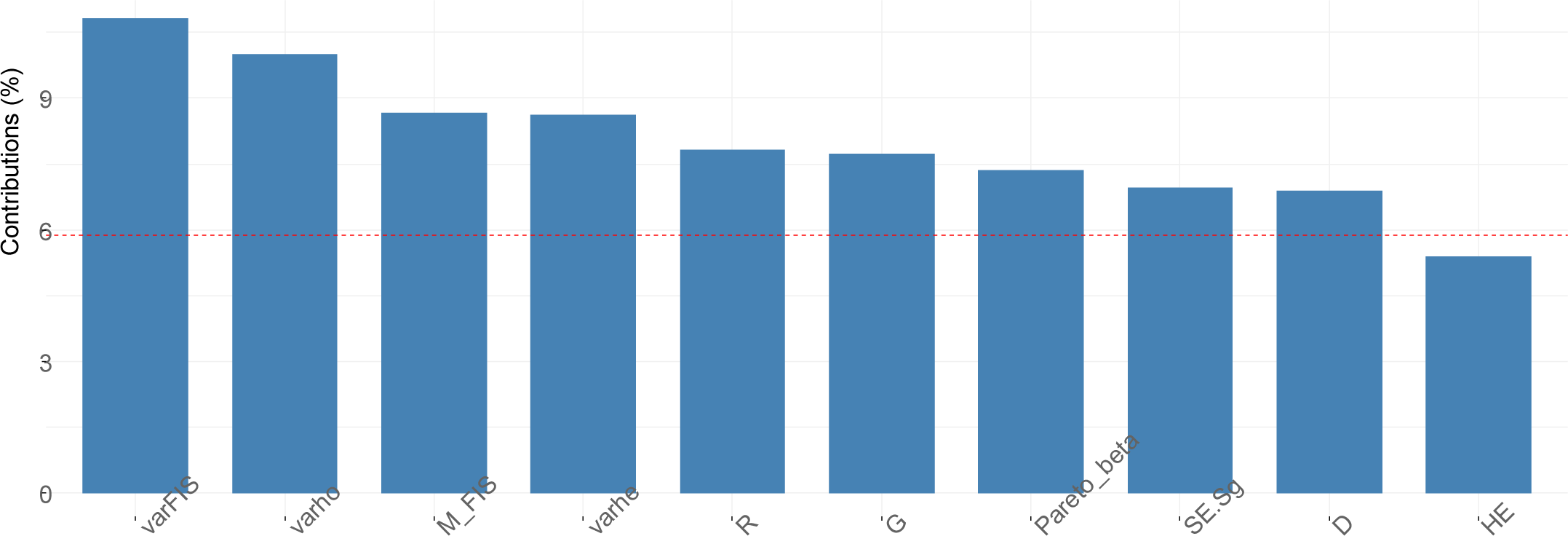
Bar plot of the genetic index contributions to the first principal component. The expected average contribution *1/(number of genetic indices)* is plotted as the red dashed line. VarFis and indices of genotypic diversity (Pareto, R, G & D) are the main contributors to the first component.

**Figure SI4.C:**
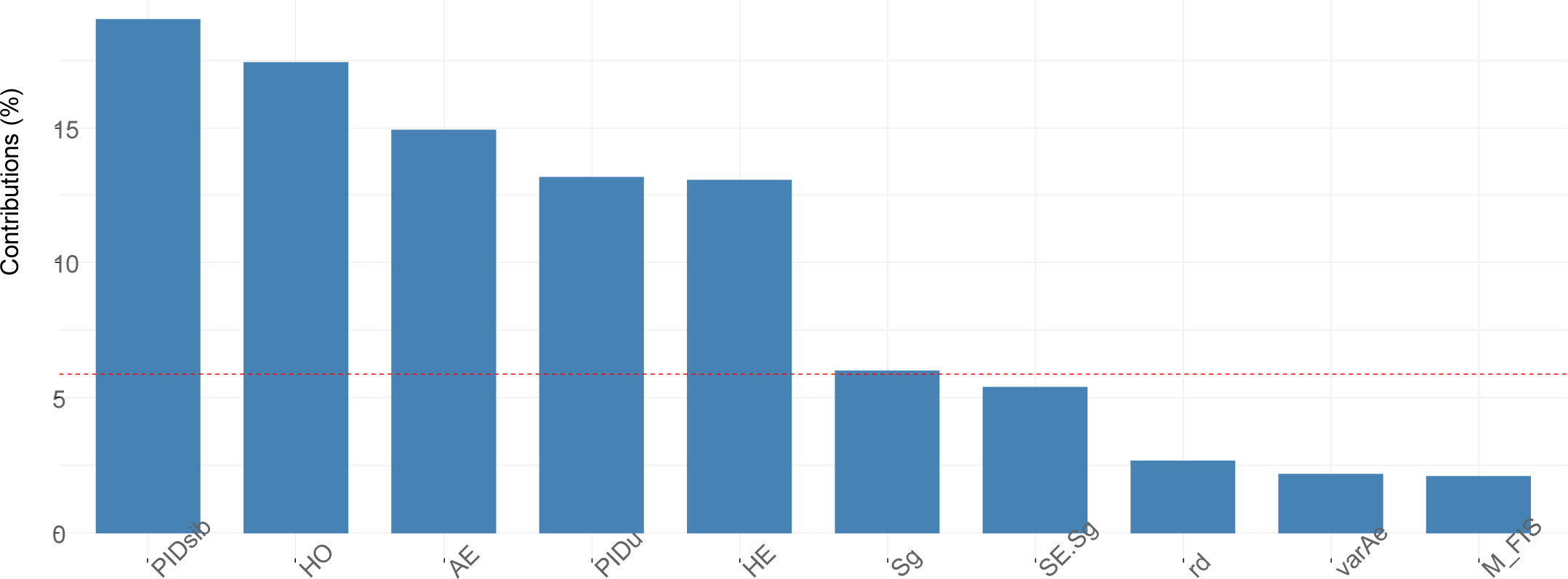
Bar plot of the genetic index contributions to the second principal component. The expected average contribution *1/(number of genetic indices)* is plotted as the red dashed line. Estimates of selfing (Sg), mean observed heterozygosity (H_O_), mean effective number of alleles (A_E_) and probabilities of identities (pid_u_ and pid_sib_) are the main contributors to the second component.

**Figure SI4.D:**
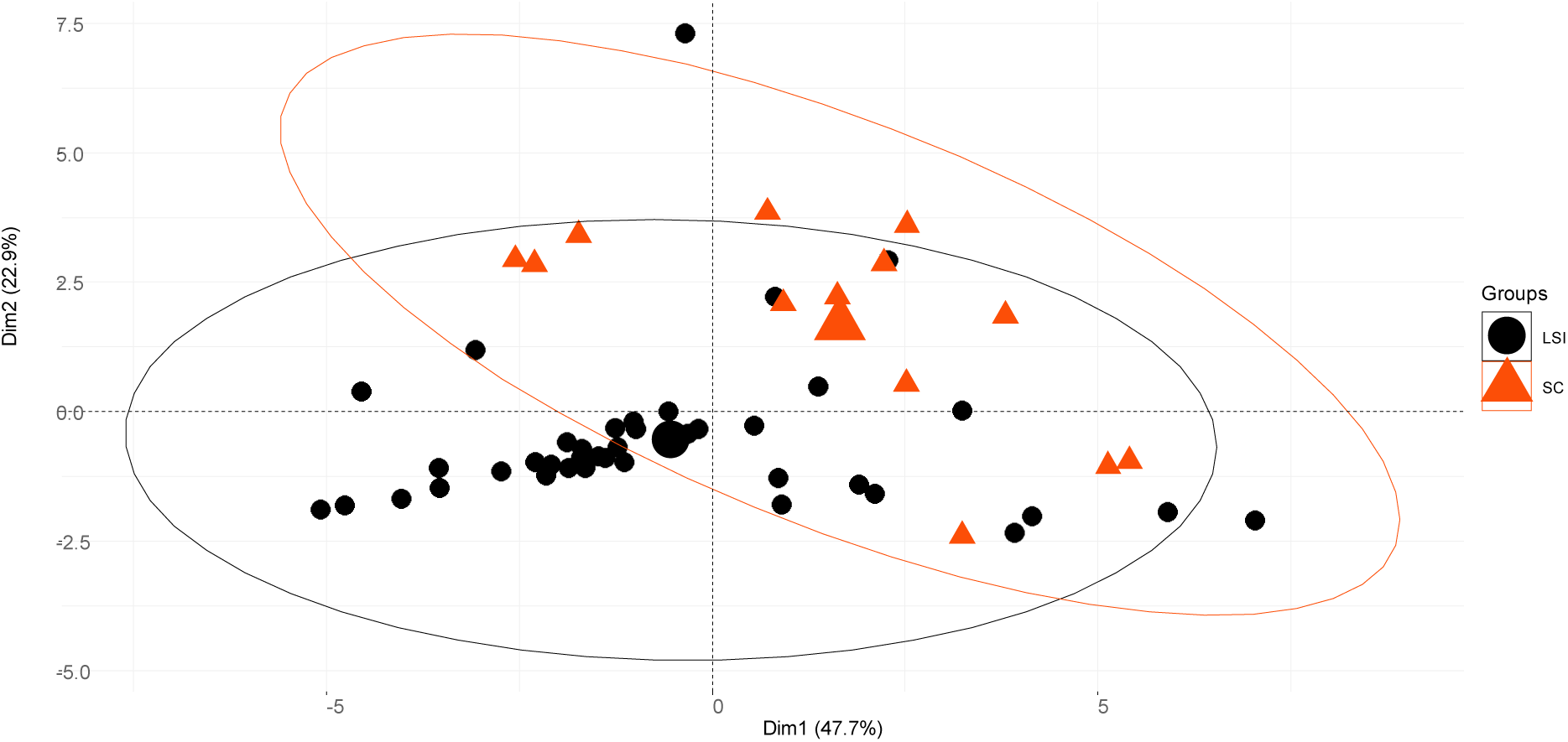
Plot of the projection of each population onto the two first principal components. SC populations are plotted as red triangles and LSI populations as black diamonds. Concentration ellipses around LSI and SC populations are plotted in black and red, respectively. Barycenters of the two groups are represented as bigger black diamond and red triangle, respectively.

**Figure SI5:**
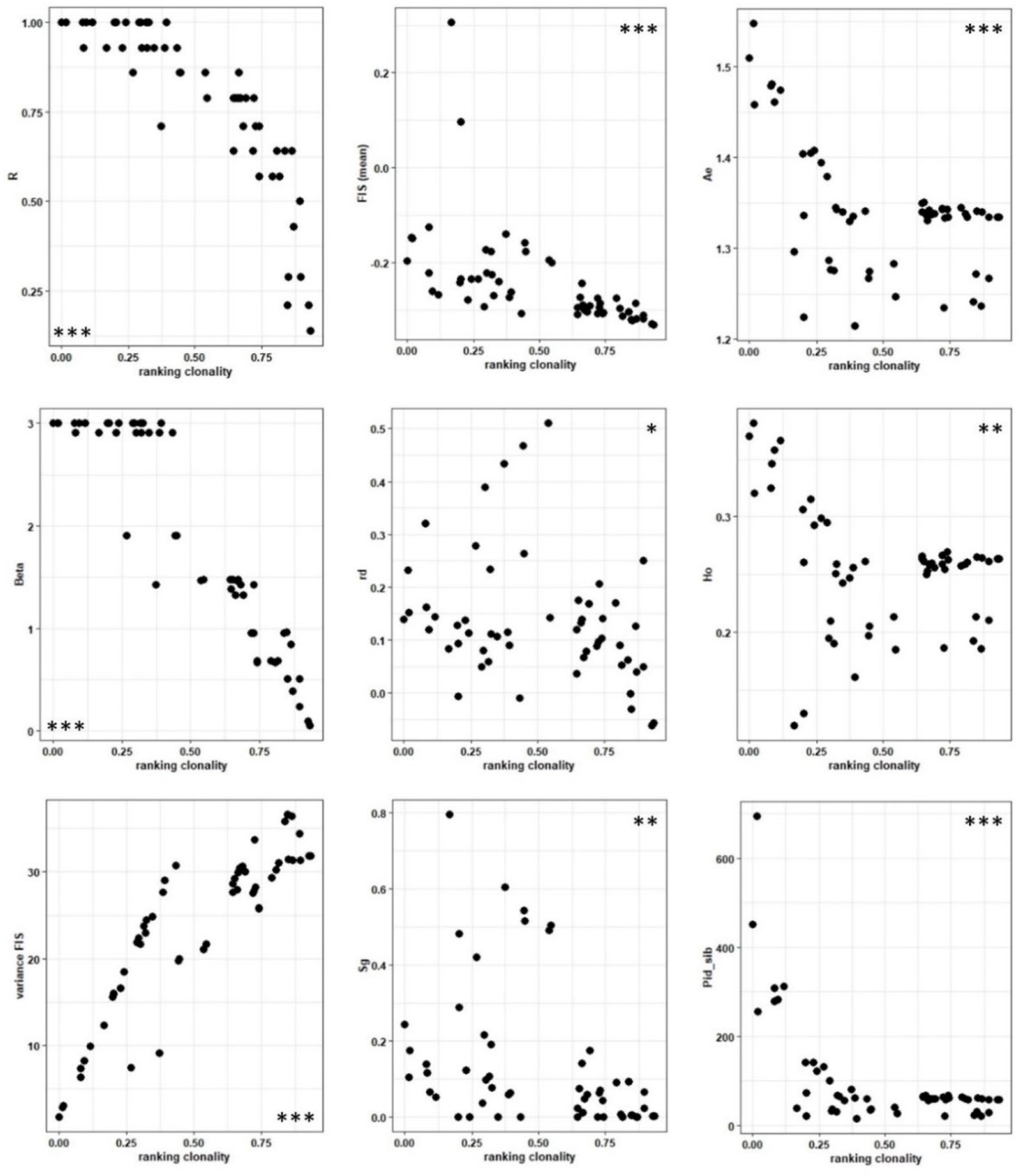
Relationship and correlation between Σ_CLON_ and genetic indices (R, Pareto β, MFis, VarFis, Ho, Ae, Sg, pidsib and *r̄_d_*). Per graph, all the 53 populations were plotted as black dots. *: p<0.05, **: p<0.01, ***: p<0.001.

**Figure SI6:**
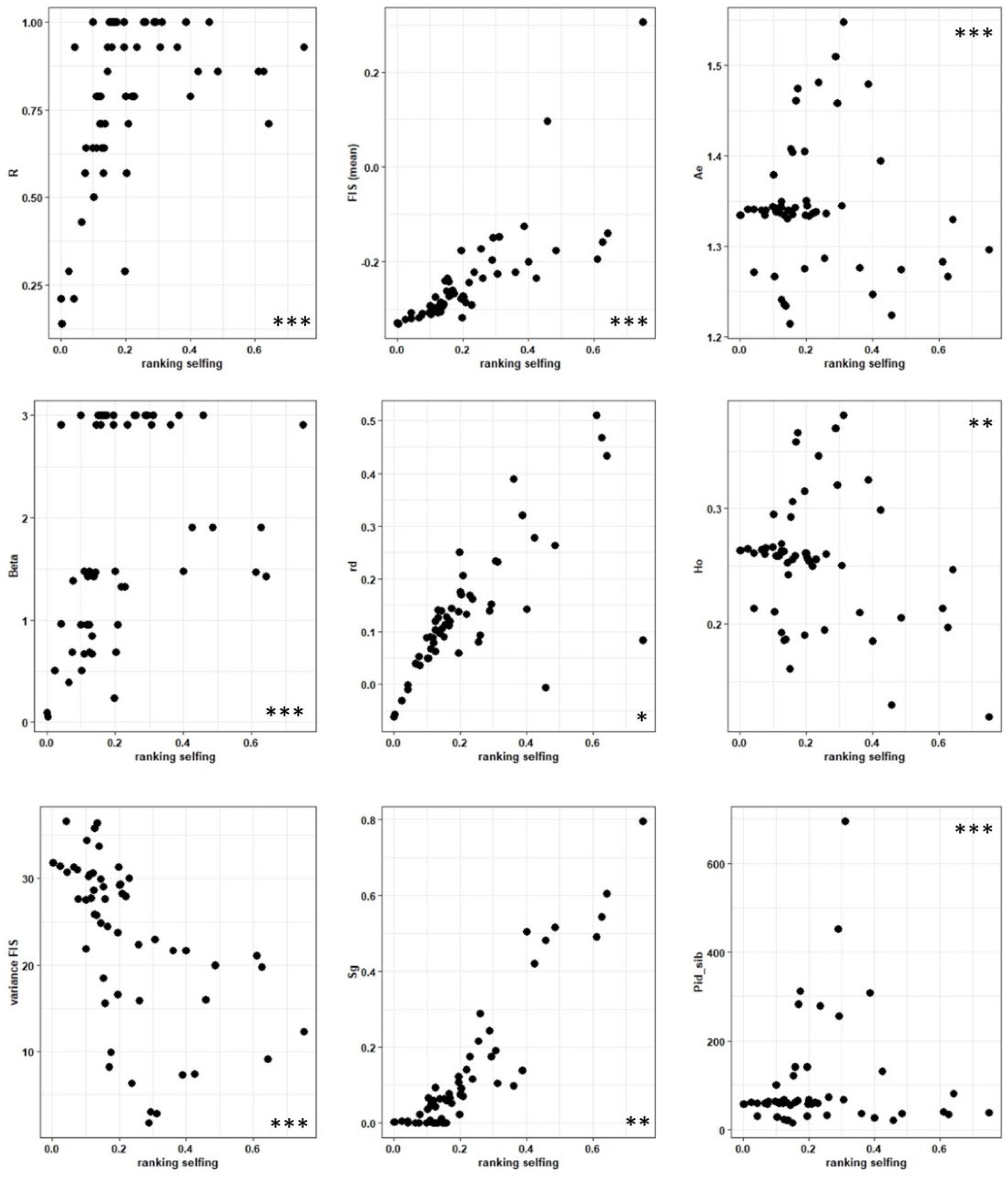
Relationship and correlation between Σ_SELF_ and genetic indices (R, Pareto β, MFis, VarFis, Ho, Ae, Sg, pidsib and *r̄_d_*). Per graph, all the 53 populations were plotted as black dots. *: p<0.05, **: p<0.01, ***: p<0.001.

**Figure SI7:**
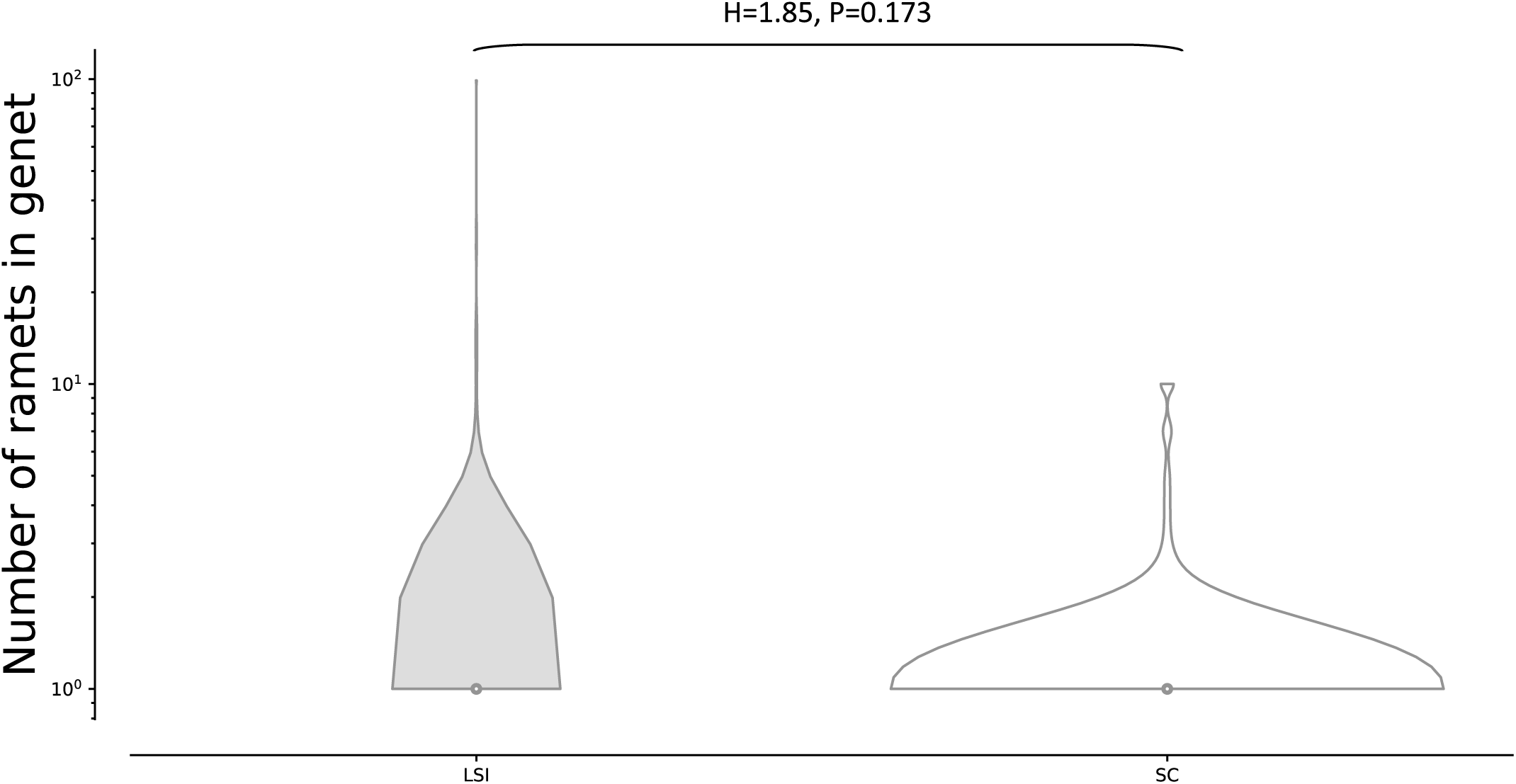
Distributions of the logarithms of the number of ramets per genet in LSI (grey) and SC (white) populations. We report the Kruskal-Wallis probability that the two distributions are identical.

## Notes

https://zenodo.org/records/10246264

### Competing Interest Statement

The authors have declared no competing interest.

### Summary of Updates

Minor changes (clarification and typos) on the third round of review.

